# Transcriptional regulation and chromatin architecture maintenance are decoupled modular functions at the *Sox2* locus

**DOI:** 10.1101/2022.02.09.479674

**Authors:** Tiegh Taylor, Natalia Sikorska, Virlana M Shchuka, Sanjay Chahar, Chenfan Ji, Neil N Macpherson, Sakthi D Moorthy, Jennifer A Mitchell, Tom Sexton

## Abstract

How distal regulatory elements control gene transcription and chromatin topology is not clearly defined, yet these processes are closely linked in lineage specification during development. Through allele-specific genome editing and chromatin interaction analyses of the *Sox2* locus in mouse embryonic stem cells, we found a striking disconnection between transcriptional control and chromatin architecture. We trace nearly all *Sox2* transcriptional activation to a small number of key transcription factor binding sites, whose deletions have no effect on promoter-enhancer interaction frequencies or topological domain organization. Local chromatin architecture maintenance, including at the topologically associating domain (TAD) boundary downstream of the *Sox2* enhancer, is widely distributed over multiple transcription factor-bound regions and maintained in a CTCF-independent manner. Furthermore, disruption of promoter-enhancer interactions by ectopic chromatin loop formation has no effect on *Sox2* expression. These findings indicate that many transcription factors are involved in modulating chromatin architecture independently of CTCF.

## INTRODUCTION

Enhancer sequences are critical regulators of gene transcription, ensuring appropriate spatiotemporal control of gene expression during development and in adult tissues (Grosveld et al., 2021). Enhancers can regulate single or multiple genes (Allahyar et al., 2018; Andersson et al., 2014; Fukaya et al., 2016; Moorthy et al., 2017), and skip over adjacent genes to regulate specific targets located at megabase-level distances away (Lettice et al., 2003; Sanyal et al., 2012). Across different tissues, chromatin modifications are more dynamic at enhancers than at promoters (Roadmap Epigenomics Consortium, 2015), and many genes are regulated by different enhancers in various cellular contexts (Bahr et al., 2018; Maqbool et al., 2020; Rodríguez-Carballo et al., 2017), suggesting that most epigenetic information is encoded at enhancers. How this regulatory complexity is encoded in the genome and how enhancers regulate the appropriate gene or genes remain open questions.

Enhancers and regulated genes are often spatially configured by chromatin-chromatin interactions, whereby gene-enhancer groups, which are linearly distant on the 2D chromosome fiber, may be brought into close proximity in 3D nuclear space (Carter et al., 2002; Palstra et al., 2003; Sanyal et al., 2012; Schoenfelder et al., 2015; Tolhuis et al., 2002). Long-range chromatin interactions are mediated by loop extrusion wherein the ring-like cohesin complex translocates bi-directionally along chromatin, bringing linearly distal regions near to one another (Davidson et al., 2019; Fudenberg et al., 2016; Li Y. et al., 2020; Sanborn et al., 2015). Chromosome conformation capture approaches have shown that the genome is partitioned into topologically associating domains (TADs) that can insulate genes in adjacent TADs from enhancer activity outside their TAD (Dixon et al., 2012; Lupiàñez et al., 2015; Sexton et al., 2012). The interaction of CCCTC-binding factor (CTCF) with the genome is enriched at TAD boundaries and the orientation of asymmetric CTCF motifs within these boundaries has a role in maintaining TAD structures (Guo et al., 2015; Nora et al., 2017; Sanborn et al., 2015; de Wit et al., 2015). CTCF has been associated in other contexts with insulator function, binding and stabilizing cohesin (Bell et al., 1999; Phillips and Corces, 2009; Wendt et al., 2008), or a means of anchoring distal enhancers to promoter-proximal CTCF-bound sites (Schuijers et al., 2018; Kubo et al., 2021). The extent to which CTCF-associated regions are required components for chromatin-chromatin contact maintenance, however, is debatable, since removal of such sites at select genomic loci has only negligible effects on chromatin topology and gene expression profiles (Despang et al., 2019; de Wit et al., 2015). Alternatively, long-range chromatin interactions can be mediated by transcription factors bound to DNA, potentially via protein dimerization events (Deng et al., 2012), clustering into nuclear hubs (Li J., et al., 2020; Mitchell and Fraser, 2008; Schoenfelder et al., 2010; Tolhuis et al., 2002) and/or formation of phase-separated condensates (Chong et al., 2018; Wei et al., 2020). Transcription factors are also able to anchor cohesin and therefore may likewise modulate loop extrusion events (Liu et al., 2021; Vos et al., 2021). Despite a clear requirement for cohesin loading/unloading dynamics in maintaining genomic architecture, perturbation studies reveal conflicting and often weak corresponding effects on the transcriptome (Liu et al., 2021; Rao et al., 2017; Schwarzer et al., 2017). Additionally, depletion of proteins involved in condensate formation at enhancers has been shown to disrupt transcription but not long-range interactions (Crump et al., 2021), demonstrating that the relationship between enhancer-promoter interactions and transcription is still not well understood.

The *Sox2* (Sex Determining Region Y-Box 2) gene encodes a transcription factor necessary for pluripotency and self-renewal in mouse embryonic stem cells (ESCs) and embryonic development (Avilion et al., 2003; Thomson et al., 2011). Deletion analyses revealed that *Sox2* transcription in mouse ESCs is regulated by the *Sox2* Control Region (SCR), a 7.3 kb cluster of transcription factor-bound regions located over 100 kb downstream of *Sox2* (Chen et al., 2012; Li et al., 2014; Zhou et al., 2014). The *Sox2* gene and the SCR are each at the border of an ESC-specific TAD that is lost upon differentiation to SOX2-dependent neural precursor cells (Bonev et al., 2017) and absent in other cell types not expressing *Sox2* (Hu et al., 2018; Stadhouders et al., 2018). Additionally, *Sox2* and the SCR appear to interact in ESCs through the formation of a chromatin loop that excludes most of the intervening DNA (Ben Zouari et al., 2020; Huang et al., 2021; de Wit et al., 2015; Zhou et al., 2014). A larger 27 kb region, comprising the SCR and two additional transcription factor-bound regions, was previously identified as a “super-enhancer” (Whyte et al., 2013), a class of genomic element originally proposed to contain multiple synergistic activators of target gene transcription. More recently, genomic interrogations of enhancer clusters have questioned the “super-enhancer” theory given that individual regions within these clusters have largely redundant functions (Hay et al., 2016; Moorthy et al., 2017). These findings are consistent with the concept of shadow enhancers, or regulatory elements that confer phenotypic robustness through their partially redundant activities (Perry et al., 2010). At the *Sox2* locus, it remains unclear how these multiple transcription factor-bound subunits within and surrounding the SCR contribute to *Sox2* transcription control or locus topology in ESCs.

To identify the sequences required for *Sox2* transcription as well as those involved in ESC-specific chromatin topology, we used allele-specific CRISPR/Cas9-mediated deletions to systematically remove all transcription factor-bound regions in the called “super-enhancer” surrounding the SCR. We find that two such regions within the SCR, *Sox2* regulatory regions (SRR) 107 and SRR111 (numerically designated according to their distance from the *Sox2* promoter), are responsible for the majority of *Sox2* transcription in ESCs; however, the deletion of these regions had no effect on interaction frequency between the SCR and the *Sox2* gene. Furthermore, removal of the sole CTCF-bound site within the SCR had no effect on either chromatin topology or *Sox2* transcription. Significant perturbation of chromatin interaction frequencies and TAD border insulation function required deletion of the entire SCR, comprising multiple transcription factor-bound sites beyond those responsible for transcriptional activation. On the other hand, insertion of CTCF motifs between *Sox2* and the SCR was able to insulate these sites from each other from a topological standpoint, but with no effect on transcription. These data show a stark uncoupling of transcription enhancement from chromatin-chromatin interaction maintenance. Furthermore, whereas enhancer function is mediated by a small number of key transcription factor-bound regions, chromatin-chromatin interaction is independent from transcriptional control and maintained in a distributed manner by many elements within the *Sox2* TAD.

## RESULTS

### Deletion of the SCR partially disrupts chromatin interactions with the *Sox2* gene in ESCs

Deletion of the SCR abrogates the majority of *Sox2* transcription in ESCs (Li et al., 2014; Zhou et al., 2014); however, the effect of SCR removal on the conformation of the locus had not been investigated. We examined the relationship between the loss of the SCR and ESC-specific chromatin architecture profile at the *Sox2* locus. To establish a locus-wide view of the chromatin contacts in both wild-type F1 ESCs (*Mus musculus*^129^ × *Mus castaneus*) and ESCs containing a homozygous deletion of the SCR (ΔSCR/ΔSCR), we subjected fixed chromatin from both lines to an allele specific 4C-seq approach (adapted from Splinter et al., 2011), using a bait region located just upstream of the SCR (**Fig 1A,B**). Regions interacting with the bait at appreciably higher levels than expected from a fitted background model (Geeven et al., 2018) were called for each biological replicate and an interaction between the SCR-proximal bait and *Sox2* gene was reproducibly identified (**Table S1**). The *Sox2*-spanning region conserved in all wild-type 4C replicates was used for quantitative interaction comparisons with other tested cells. Relative to wild-type cells, ΔSCR/ΔSCR cells showed a significant 28% decrease (*P* = 0.02) in relative contact frequency (**Fig 1C**). This finding suggested that regions within the SCR contribute to the maintenance of ESC-specific genomic configurations at the *Sox2* locus. An alternative mechanistic explanation, however, was also possible: ESCs containing a homozygous deletion of the SCR are partially differentiated and exhibit markedly reduced SOX2 protein levels (Zhou et al., 2014). We therefore raised the question of whether the observed reduction in SCR-*Sox2* gene interaction frequency in these cells was mediated by *trans* mechanisms associated with the depletion of SOX2 protein, which might be required to anchor these chromatin contacts, rather than by deletion of the *Sox2* enhancer DNA *in cis*. We assessed allele-specific contact frequencies across the *Sox2* locus in cells carrying a heterozygous deletion of the SCR, which contained wild-type equivalent SOX2 protein levels (Zhou et al., 2014). The allele containing an intact SCR (**Fig 1C**, WT allele) exhibited a chromatin contact profile that mirrored that observed in wild-type cells (*P* = 0.46), whereas the relative *Sox2*-SCR contact frequency of the allele with the deletion was reduced by 24% (ΔSCR allele; *P* = 0.04) compared to wild-type levels. This loss is not significantly different to the reduction observed in cells containing homozygous deletion of the SCR (*P* = 0.74) (**Fig 1C**). These results thus indicate that the loss of the SCR *in cis* directly accounts for reduced interactions within the *Sox2* locus. Interestingly, the majority of the interaction is apparently maintained with complete loss of the SCR, which confers more than 80% of *Sox2* transcriptional activity. These data suggest that genome architecture and expression control may be decoupled at this locus, and that additional *cis*-elements are required to maintain local chromatin structure.

**Figure 1.**
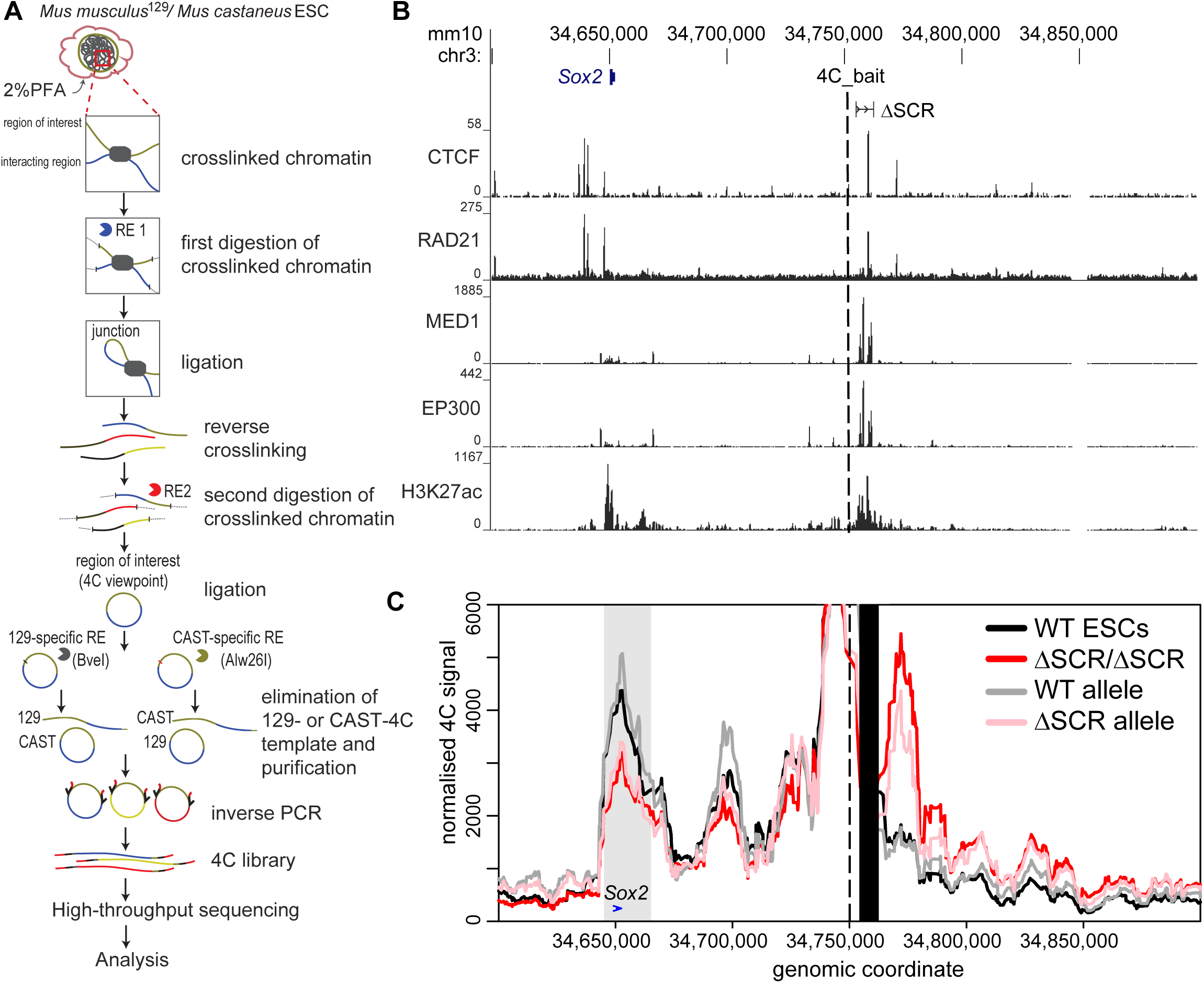
Deletion of the SCR partially disturbs chromatin interactions with the *Sox2* gene in ESCs. A) Schematic of the allele-specific 4C approach. Restriction Enzyme (RE), paraformaldehyde (PFA). B) The region surrounding the *Sox2* gene is displayed on the UCSC Genome Browser (mm10). The SCR deletion (ΔSCR) is shown and the 4C bait region is indicated as a dashed line. ChIP-seq conducted in ESCs is shown below for CTCF, RAD21, MED1, EP300 and H3K27ac. C) 4C data are shown for wild-type (WT, black, n=4), homozygous ΔSCR/ΔSCR cells (red, n=4), and heterozygous ΔSCR cells. Data from the heterozygous cells are displayed separately for the WT (grey, n=3) and ΔSCR alleles (pink, n=4). The dashed line indicates the location of the 4C bait region. The black box indicates the deleted region. The grey box indicates the bait-interacting region surrounding the *Sox2* gene (blue arrow). Compared to WT cells, a significant decrease in relative interaction frequency of the 4C bait region with the *Sox2* gene was observed for homozygous ΔSCR/ΔSCR cells (*P* = 0.02), and the ΔSCR allele in heterozygous ΔSCR cells (*P* = 0.04), but not the WT allele in heterozygous ΔSCR cells (*P* = 0.46).

### Two transcription factor-bound regions are jointly responsible for SCR-mediated enhancement of *Sox2* transcription

We next sought to assess which sub-regions within the SCR contribute to *Sox2* transcription. We previously established that, of the four transcription factor-bound regions within the SCR (SRRs 106, 107, 109 and 111, **Fig 2A**), only SRR107 and SRR111 are capable of upregulating transcription of a reporter gene in ESCs (Zhou et al., 2014). To examine whether the same holds true in a genomic context, we created ESC clones with heterozygous deletions of either SCR sub-region on the 129 allele (**Fig S1** and **Tables S2 and S3**). Allele-specific *Sox2* transcript level quantification analysis (Moorthy and Mitchell, 2016) allowed us to assess the endogenous activation potential of either region. We observed that the loss of SRR107 from the 129 allele is accompanied by a modest, but significant alteration in the allelic ratio of *Sox2* transcripts, with a 27% reduction in transcript levels from the allele carrying the deletion. Removal of SRR111 caused a weaker (14%), non-significant reduction in *Sox2* transcript production from the 129 allele (**Fig 2B**). A compound deletion made by deleting SRR111 on the 129 allele in a genomic background already lacking SRR107 on the same allele demonstrated a much larger (70%) decrease in allele-specific *Sox2* expression (ΔSRR107+111/+, **Fig 2B**). Furthermore, this reduction in transcript abundance was not statistically different from that observed in clones lacking the entire SCR. Notably, the allelic imbalance of *Sox2* transcription was essentially the same whether or not the intervening region between SRR107 and SRR111 was also deleted, as indicated by the expression results for ΔSRR107-111/+ and ΔSRR107+111/+ cells (**Fig 2B**). These data indicate that SRR107 and SRR111, acting in a partially redundant manner, underlie the transcriptional regulatory power of the SCR in coordinating ESC-specific *Sox2* transcription.

**Figure 2.**
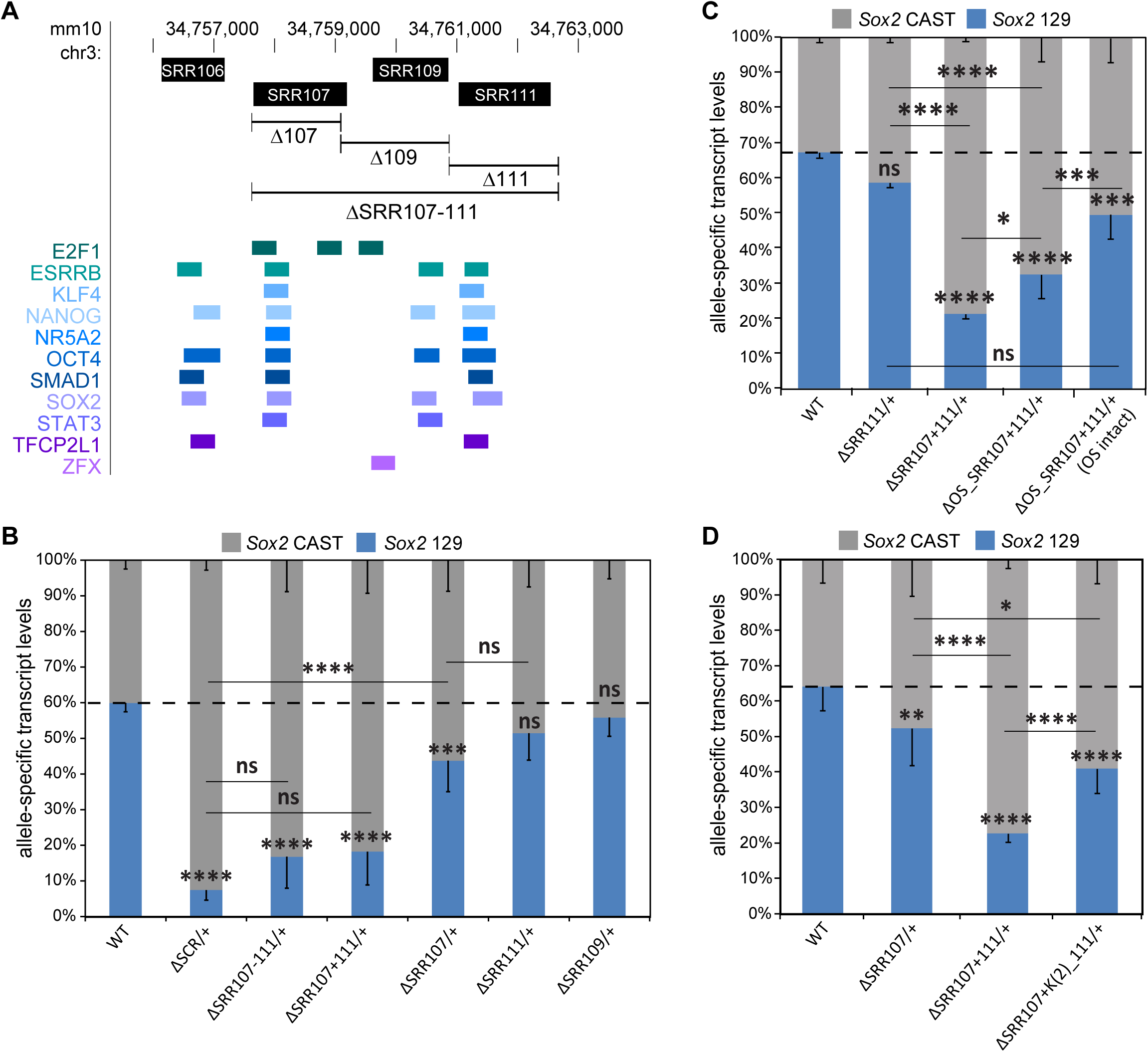
SRR107 and SRR111 have a partially redundant role and are required for the transcription-enhancing capacity of the SCR. A) The SCR genomic region is displayed on the UCSC Genome Browser (mm10). *Sox2* regulatory regions (SRR, above) correspond to transcription factor-bound regions derived from ESC ChIP-seq datasets compiled in the CODEX database (below). In B-D, allele-specific primers detect *Sox2* musculus (129; blue) or castaneus (CAST; grey) mRNA by RT-qPCR from F1 ESC clones from the indicated genotype. Expression levels for each allele are shown relative to the total transcript levels. Error bars represent SD, n ≥ 3. Significant differences from the WT values are indicated: (*) *P* < 0.05, (**) *P* < 0.01, (***) *P* < 0.001, (****) *P* < 0.0001, ns = not significant. B) Deletion of both SRR107 and SRR111 (ΔSRR107-111/+; ΔSRR107+111/+) causes a similar reduction in *Sox2* transcript levels as for deletion of the entire SCR. C) Deletion of the OCT4:SOX2 (OS) motif in SRR107 (ΔOS_SRR107+111/+) reduces *Sox2* transcript levels on the linked allele. Clones with nucleotide deletions near but not disrupting the OS motif [ΔOS_SRR107+111/+ (OS intact)] displayed increased transcription of *Sox2* on the linked allele compared to clones with a deleted OS motif. D) Deletion of two KLF4 motifs in SRR111 [ΔSRR107+ K(2)_111/+] reduces *Sox2* transcript levels on the linked allele.

Since *Sox2* transcription was shown to be significantly reduced upon deletion of both SRR107 and SRR111 on the same allele, these enhancers were investigated for the presence of core ESC transcription factor binding motifs that may promote the activity of these regions. Using the JASPAR GeneReg Database tool (Sandelin et al., 2004), we uncovered the presence of multiple high-scoring transcription factor motifs, including a POU5F1:SOX2 composite motif in SRR107 and two KLF4 motifs in SRR111 (**Fig S2A**). Overlapping of ChIP-seq datasets extracted from the CODEX database (Sánchez-Castillo et al., 2015) confirmed the association of the corresponding transcription factor proteins over these sites (**Fig 2A**). Because of the partially redundant functions of the two enhancers, we targeted each of these motifs for removal in clones carrying only one intact enhancer (in which the motif of interest resided) with the other SRR deleted in the same allele. Clones carrying micro-deletions of the targeted motifs (**Fig S2B**) were subjected to allele-specific expression analyses. We found that deletion of the high-scoring POU5F1:SOX2 motif in SRR107 resulted in a transcript reduction level close to that of the loss of the entire SRR107 in an SRR111 deletion-carrying background (**Fig 2C**). Importantly, clones carrying slightly off-target micro-deletions that retained an intact POU5F1:SOX2 motif do not show as great a decrease in allele-specific *Sox2* transcript levels. Similarly, the loss of both KLF4 motifs in SRR111 also caused a significant reduction of this region’s activity in an SRR107 deletion-carrying background (**Fig 2D**). These results suggest that deletion of the designated motifs accounts for the reduced expression phenotype we observed upon deletion of the entire transcription factor-bound sub-region. Collectively, these findings support a model for a distal gene regulation mechanism controlling *Sox2* transcription in ESCs that depends upon ESC-specific transcription factors bound at these two regions.

### Enhancer activity and CTCF association are dispensable for distal chromatin contacts within the *Sox2* locus

Aside from SRR107 and SRR111, which jointly drive the transcriptional enhancer activity for *Sox2*, the SCR also contains a prominent CTCF ChIP-seq peak at SRR109 (**Fig 1B**). CTCF has been associated with anchoring distal enhancers to promoter-proximal CTCF-bound sites (Kubo et al., 2021); however, a previous study identified only a slight decrease in the observed interaction frequency of the SCR with *Sox2* after bi-allelic removal of the core 16 bp within the CTCF motif in the SRR109 region (de Wit et al., 2015). To assess whether SRR109 might function as an intra-SCR loop anchor, we generated cell lines containing a heterozygous deletion of SRR109 on the 129 allele (**Fig S3**) and subjected these clones to allele-specific 4C-seq and expression analysis. In line with previous perturbations of the SRR109 CTCF site (de Wit et al., 2015) and enhancer reporter assays (Zhou et al., 2014), heterozygous SRR109 deletion had only negligible effects on the allelic balance of *Sox2* expression levels (**Fig 2B**), indicating that this region has next to no direct enhancer activity in ESCs. Moreover, we observed no significant differences in 129 allele-derived chromatin-chromatin contact profiles between cells lacking one copy of SRR109 and wild-type cells (*P* = 0.4, **Fig 3A**), suggesting that this CTCF-bound element is not required for the genomic proximity between *Sox2* and the SCR. The observation that SRR107 and SRR111 are required for maintenance of *Sox2* transcription in ESCs led us to hypothesize that these two regions are combinatorially responsible for anchoring chromatin interactions between the SCR-proximal region and the *Sox2* gene. Allele-specific 4C analysis of cells lacking both SRR107 and SRR111, however, revealed no significant differences in contact frequencies between the distal SCR-proximal region and *Sox2* in ΔSRR107+111/+ cells and wild-type cells (P = 0.6, **Fig 3B**). These findings support the notion that both SRR107 and SRR111 appear to be dispensable for the SCR-*Sox2* interaction.

**Figure 3.**
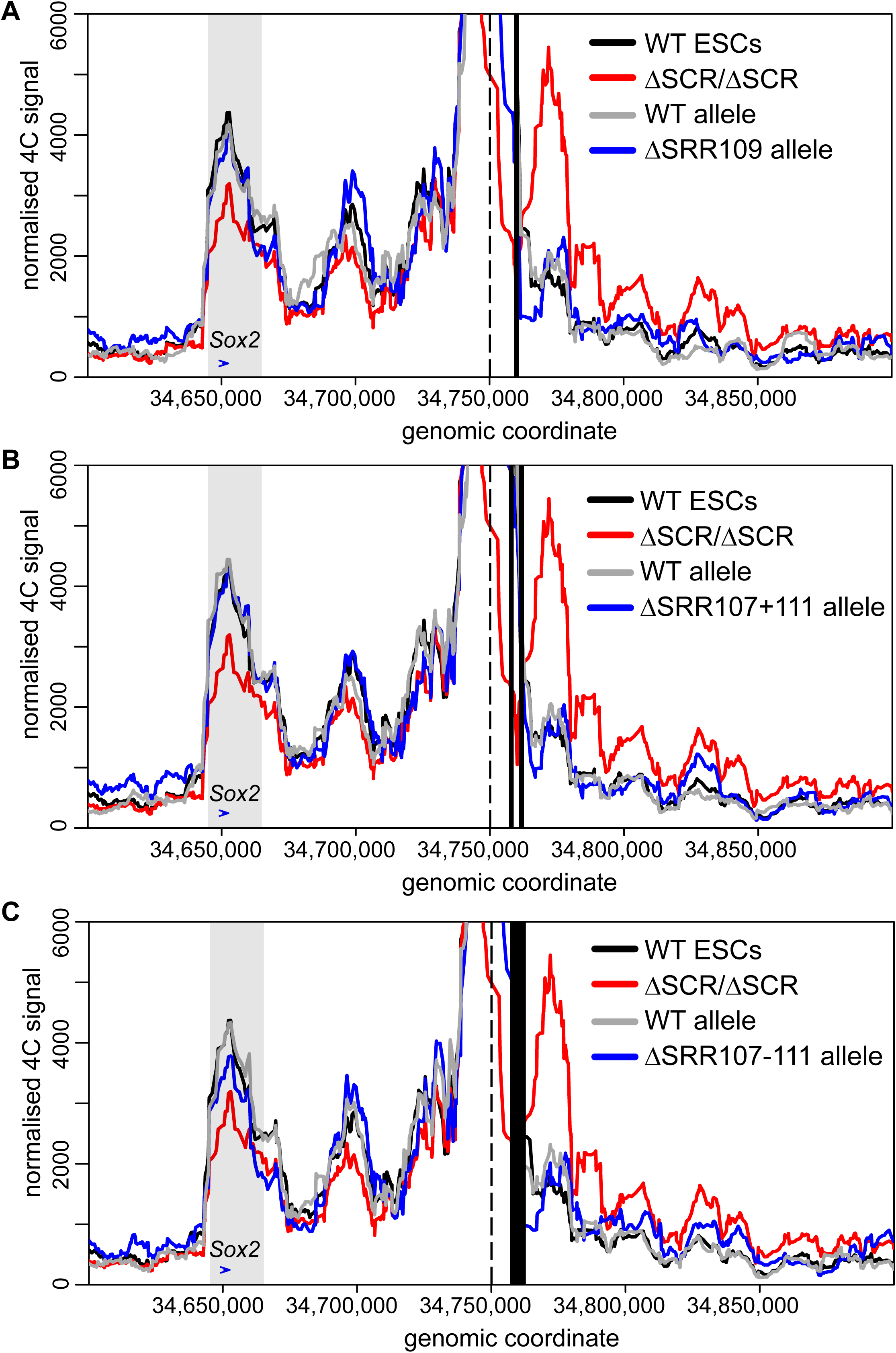
The SRR107 and SRR111 enhancers and the SRR109 CTCF-bound region are dispensable for the interaction between the SCR proximal region and the *Sox2* gene. 4C data are shown for wild-type (WT, black, n=4), homozygous ΔSCR/ΔSCR cells (red, n=4), and heterozygous ΔSRR cells (grey/blue, n=2 for each allele). Heterozygous deletions of SRR109 (A), SRR107 and SRR111 (B), or SRR107 to SRR111 (C) are shown in blue, with the WT allele shown in grey. In each figure the dashed line indicates the location of the 4C bait region. The black box indicates the deleted region(s). The grey box indicates the bait-interacting region surrounding the *Sox2* gene (blue arrow), which is not significantly altered upon deletion of SRR109 (*P* = 0.4), SRR107 and SRR111 (*P* = 0.6), or SRR107 to SRR111 (*P* = 0.08).

We next considered that SRR109 might cooperate with the two enhancer components of the SCR to support its interaction with the *Sox2* promoter. To test this hypothesis, we used ΔSRR107-111/+ ESCs, which lacked both enhancers and the CTCF-bound region on the same allele (**Fig S3**). Here, we did observe a reduction (17%) in the chromatin interaction profile between the SCR-proximal region and the *Sox2* gene on the SRR107-SRR111 deletion-carrying allele compared to wild-type cells. However, this observation did not meet the critical value for statistical significance in our analysis (*P* = 0.08, **Fig 3C**). This finding suggests that SRRs 107, 109, and 111 may minimally support the interaction of the SCR with the *Sox2* gene, although other genetic elements likely contribute to the bulk (∼83%) of the interaction. Overall, our results indicate a striking decoupling at this locus between genetic elements responsible for transcriptional activation (key transcription factor motifs within SRR107 and SRR111) and those influencing chromatin architecture, where the sole CTCF-bound site within the SCR appears to have only a minor contribution.

### A downstream CTCF-bound region is not responsible for the remaining *Sox2-*SCR-proximal region interaction in SCR deletion-carrying cells

Since both homozygous and heterozygous SCR deletions were not sufficient to abolish chromatin interactions between the SCR-proximal region and the *Sox2* gene, we searched for other candidate regions surrounding the SCR that might support the bulk of the remaining interactions. We postulated that a CTCF-bound region (distal CTCF, dCTCF in **Fig 4A**) downstream of the SCR may be involved in stabilizing the interaction between the SCR-proximal region and *Sox2* when the SCR is deleted. Co-bound by CTCF and RAD21 (**Fig 1B**), this site could act as a chromatin contact-anchoring region. Furthermore, given the functional redundancy between enhancer regions regulating *Sox2* transcription, it is plausible that CTCF-bound regions may also act redundantly to stabilize chromatin interactions. The dCTCF motif is oriented away from *Sox2* and not expected to participate in a direct interaction with the gene via stalled loop extrusion intermediates; however, other *in silico* data suggests that divergent CTCF pairing (such as that between SRR109 and dCTCF) cooperates to reinforce TAD and intra-TAD loops (Nanni et al., 2020). To evaluate this possibility, we created additional deletions at regions outside the SCR (**Fig S4A**). We extended the distal deletion to include the SCR and the downstream CTCF-bound region (ΔSCR-dCTCF/+), thus removing both CTCF-bound sites. However, these deletions demonstrated only a marginally increased loss of chromatin interaction (36%) compared to that observed with deletion of the SCR alone (28%), a variation in interaction frequencies that was not significantly different (*P* = 0.5, **Fig 4B**).

**Figure 4.**
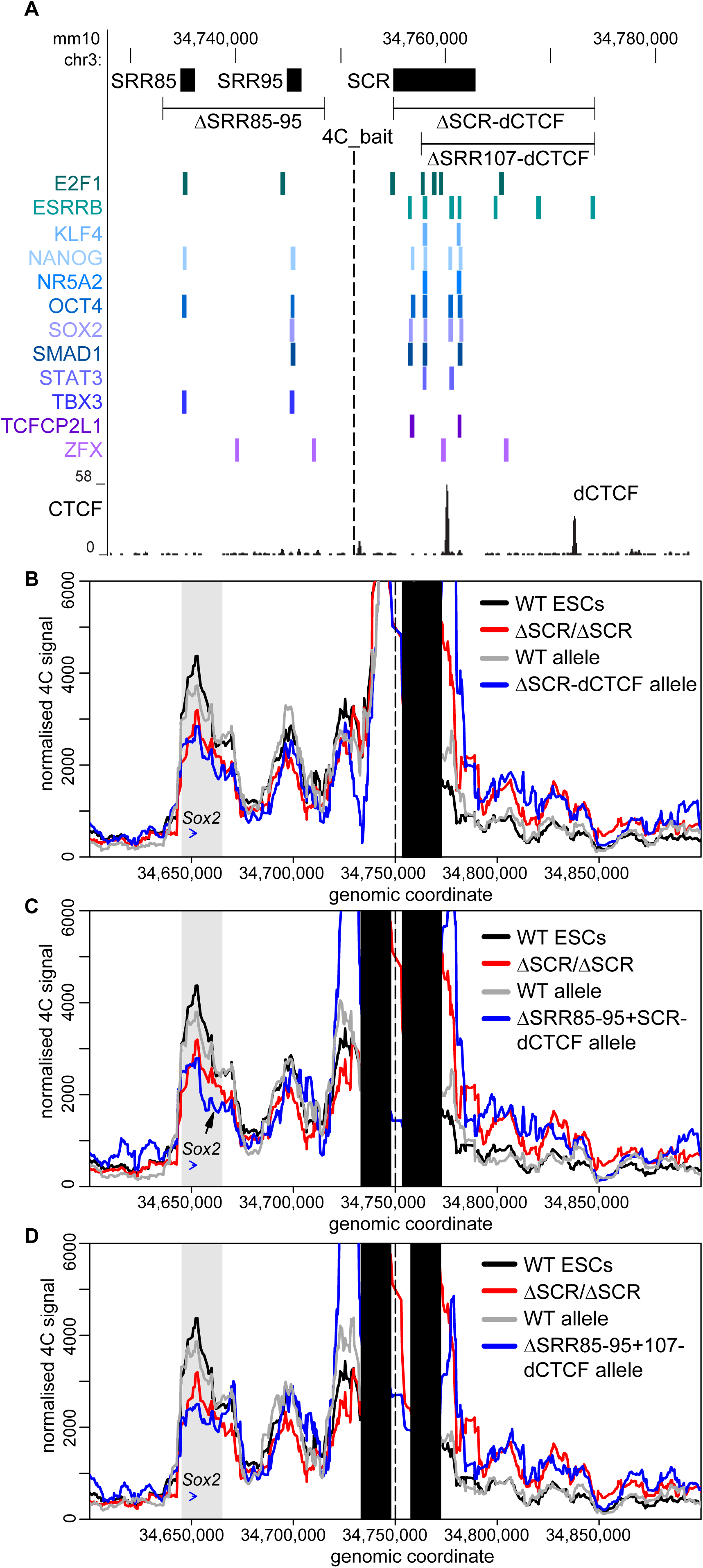
Additional CTCF- and transcription factor-bound regions surrounding the SCR support the interaction between the SCR proximal region and the *Sox2* gene. A) The transcription factor-bound regions surrounding the SCR is displayed on the UCSC Genome Browser (mm10). *Sox2* regulatory regions (SRR) and the SCR (above) correspond to transcription factor-bound regions derived from ESC ChIP-seq datasets compiled in the CODEX database (below). CTCF ChIP-seq conducted in ESCs is shown (below) and the distal CTCF (dCTCF)-bound region is marked. In B-D, 4C data are shown for wild-type (WT, black, n=4), homozygous ΔSCR/ΔSCR cells (red, n=4), and heterozygous deletion-carrying cells (grey/blue, n=2 for each allele). Heterozygous deletions of the SCR to dCTCF (B), SRR85 to SRR95 and SCR to dCTCF (C), or SRR85 to SRR95 and SRR107 to dCTCF (D) are shown in blue, with the WT allele shown in grey. In each figure, the dashed line indicates the location of the 4C bait region. The black box indicates the deleted region(s). The grey box indicates the bait-interacting region surrounding the *Sox2* gene (blue arrow). Compared to WT cells, a significant decrease in relative interaction frequency of the 4C bait region with the *Sox2* gene was observed for ΔSCR­dCTCF/+ *(P =* 0.006), ΔSRR85-95+SCR-dCTCF/+ *(P =* 0.005), or ΔSRR85-95+107-dCTCF/+ *(P =* 0.01) cells. The arrow indicates a loss of interaction downstream of the *Sox2* gene after deletion of ΔSRR85-95+ SCR-dCTCF/+.

Upstream of the SCR, the locus contains two separate transcription factor-bound regions located 85 and 95 kb downstream of the *Sox2* promoter (SRRs 85 and 95; **Fig 4A**). These regions were previously shown to lack enhancer activity in a reporter assay (Zhou et al., 2014). Their deletion, either on their own (ΔSRR85-95/+) or in combination with the SCR(ΔSRR85-95+SCR-dCTCF/+), revealed that SRR85 and SRR95 also lack enhancer activity in their endogenous genomic context (**Fig S4B**). We tested whether these elements may play more of an architectural role in stabilizing a transcription factor-bound “hub” instead. Upon producing the compound deletion ΔSRR85-95+SCR-dCTCF/+, which removes all major CTCF and transcription factor-bound regions in and around the SCR, we surprisingly noted only a subtle further reduction (36%) in interaction frequency compared to deletion of the SCR alone (28%, **Fig 4C**). The extent of interaction loss appears identical to that caused by deletion of SCR to dCTCF alone, which would suggest that SRR85 and SRR95 play no additional role in chromatin architecture. However, we observed a reduction of interactions with regions just downstream of *Sox2* in ESCs lacking the SRR85 to SRR95 region (**Fig 4C**; arrow).

When comparing all the locus deletions discussed thus far, we noted that the significant reduction in chromatin contacts at the *Sox2* locus was only achieved by deletion of the entire SCR; the combined deletion of SRR109 (and its resident CTCF site) and the two principal enhancer elements (ΔSRR107-111/+, **Fig 3**) caused only a minimal disruption in the interaction frequencies with the *Sox2* gene. We thus considered a possibility that the one remaining transcription factor-bound region in the SCR, SRR106, could safeguard the local chromatin architecture at the locus. To test this possibility, we generated an ESC clone combining deletions of all of the previously interrogated regions on the 129 allele but truncating the SCR to dCTCF deletion to leave SRR106 intact (ΔSRR85-95+107-dCTCF/+, **Fig 4D**). Distal interaction frequency with *Sox2* decreased by 43% in ΔSRR85-95+107-dCTCF/+ cells compared to wild type (*P =* 0.01). Yet, this chromatin contact profile is quantitatively similar and not different statistically from the reduction caused by deletion of the SCR (*P* = 0.3, **Fig 4D**). Thus SRR106 does not appear to support the maintenance of local chromatin topology in the absence of other transcription factor-bound regions. Overall, these results reinforce the notion of a decoupling of *Sox2* transcriptional control and chromatin architecture within the locus. Whereas two single enhancer elements confer the vast majority of transcriptional control in pluripotent ESCs (at least under the conditions of the experiment), chromatin architecture supporting interactions between SCR-proximal regions and the *Sox2* gene is widely distributed over many contributing elements. Much of the chromatin contact profile (>50% of wild-type levels) was maintained in the absence of *Sox2* transcription and persisted even in cells with the most extreme deletions generated during this study (**Table S1**).

### The downstream *Sox2* TAD border is insulated by the entire SCR in a CTCF-independent manner

Publicly available Hi-C data have shown that the *Sox2* gene and the SCR each reside near a TAD boundary thereby restricting interactions with flanking chromatin outside the ESC-specific *Sox2* TAD (**Fig 5A** and **Fig S5A**). When we evaluated interaction frequencies between the SCR-proximal bait region and genomic regions outside its resident TAD, we noted that removal of the SCR did not generate any specific ectopic interactions up- or downstream of the TAD borders, but did cause a general increase in 4C signal downstream of the SCR (**Fig 1C**). Extending the view of the 4C results to a larger window, it was clear by visual inspection that the SCR deletion caused a general increase in basal interaction frequency with the entire downstream chromatin up to the next TAD border, whereas interaction frequencies with the upstream TAD were unaffected (**Fig 5B**). Although this type of boundary function is often associated with CTCF binding, we noted that removal of SRR109, which is the only region bound by CTCF within the SCR, neither disrupts the SCR boundary nor causes any change in the contact profiles with either the upstream or downstream TADs. We next analyzed the interaction frequencies across the entire downstream TAD segment and the genomic region of the same size (325 kb) in the upstream TAD in ESCs harboring deletions of regions we had prioritized as candidate regulators of chromatin topology at the *Sox2* locus. We observed that only removal of the entire SCR, either alone or in combination with deletions of SRR85 and SRR95, and/or dCTCF, were associated with the observed “leakiness” of the downstream TAD border (**Fig 5C**). This finding indicates that, similar to the *Sox2*-SCR interaction, the TAD boundary is robustly maintained by multiple genomic elements acting in a partially redundant manner and is not conferred by the CTCF site alone.

**Figure 5.**
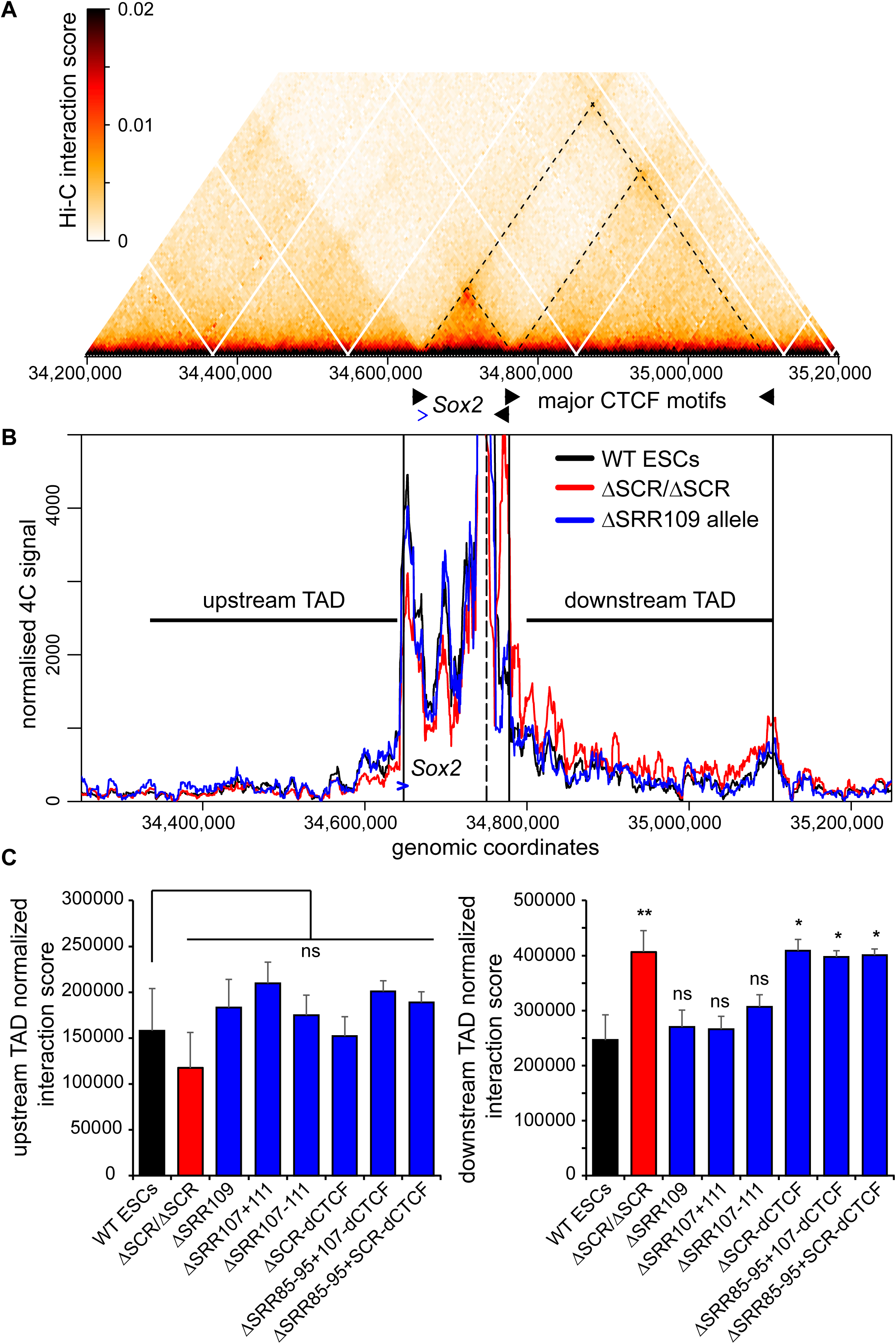
SCR deletion affects the *Sox2*-SCR TAD boundary and causes increased interaction with the downstream TAD. A) Hi-C data from ESCs (acquired from Bonev et al., 2017) indicating the frequency of occurring interactions surrounding Sox2. The dashed lines correspond to the TAD boundaries at the *Sox2* promoter, SCR and the downstream TAD boundary. B) 4C data is shown for wild-type (WT, black, n=4), homozygous ΔSCR/ΔSCR cells (red, n=4), and the ΔSRR109 allele in heterozygous ΔSRR109/+ cells (blue, n=2). The dashed line indicates the location of the 4C bait region. The vertical black lines indicate TAD boundaries shown in A. The *Sox2* gene is indicated by a blue arrow. The horizontal lines indicate the 325 kb upstream and downstream TAD regions used to calculate the interaction score shown in C. C) Normalized interaction scores for the downstream (left) and upstream (right) TAD regions are shown for the indicated F1 ESC clones, revealing that only the full SCR deletion significantly increases interaction frequencies between the SCR proximal region with the downstream TAD. The blue bars mark the interactions observed for the deleted allele in the indicated heterozygous deletion-carrying clones. Significant differences from the interaction in WT cells are indicated by * (*P* < 0.05), ** (*P* < 0.01), ns = not significant. Data shown are an average of 2-4 biological replicates, with error bars representing SD.

Since the CTCF binding sites within the ESC *Sox2* TAD examined in this study appeared to be completely dispensable for local chromatin topology, we next asked whether the CTCF protein was required for *Sox2* expression and *Sox2*-SCR interaction. We re-analyzed Hi-C data from ESCs before and after acute CTCF depletion via an engineered auxin-inducible degron, where large-scale disruption in TADs had previously been reported. Notably, RNA-seq showed only minor transcriptomic changes, including no effect on *Sox2* expression (Nora et al., 2017). In line with our own findings, CTCF ablation caused negligible (<1.04-fold) changes to the *Sox2*­SCR interaction frequency or overall *Sox2* TAD structure (**Fig S5B**), suggesting that local chromatin architecture is indeed CTCF-independent.

### *Sox2* transcription is maintained despite perturbation of chromatin contacts with the SCR

The deletion experiments described thus far have allowed for the fine functional dissection of the SCR and SCR-proximal elements to evaluate the role these regions have in regulating *Sox2* expression and locus topology; however, these data did not indicate whether *Sox2* TAD architecture is in fact important for transcriptional control. All ESC lines harboring deletions that perturbed chromatin interactions had removed the SRR107 and SRR111 elements required for efficient *Sox2* expression. To assess the functional significance of the chromatin interactions at the *Sox2* locus, we chose to disrupt these endogenous *Sox2*-SCR interactions while keeping the SCR intact. We engineered an FRT/F3 cassette at a site located between the *Sox2* gene and the SCR in the 129 allele of F1 ESCs (**Figs S6** and **S7** and **Table S4**). We were thus able to site-specifically insert putative insulator sequences of interest by recombinase-mediated cassette exchange and assess their effect on allele-specific chromatin topology and *Sox2* expression levels (**Fig 6A**). This approach was previously applied to the *Sox2* locus; Huang et al. reported that efficient transcriptional insulation (i.e. reduction of *Sox2* expression levels by ∼30-40%) required the insertion of tandem copies of CTCF motifs and flanking sequences (comprising ∼4 kb total inserted sequence) to alter endogenous *Sox2*-SCR interactions (Huang et al., 2021). However, this study only assessed chromatin topology for insertions where strong transcriptional inhibition had been observed; whether structural perturbations could occur in the absence of transcription had not been determined.

**Figure 6.**
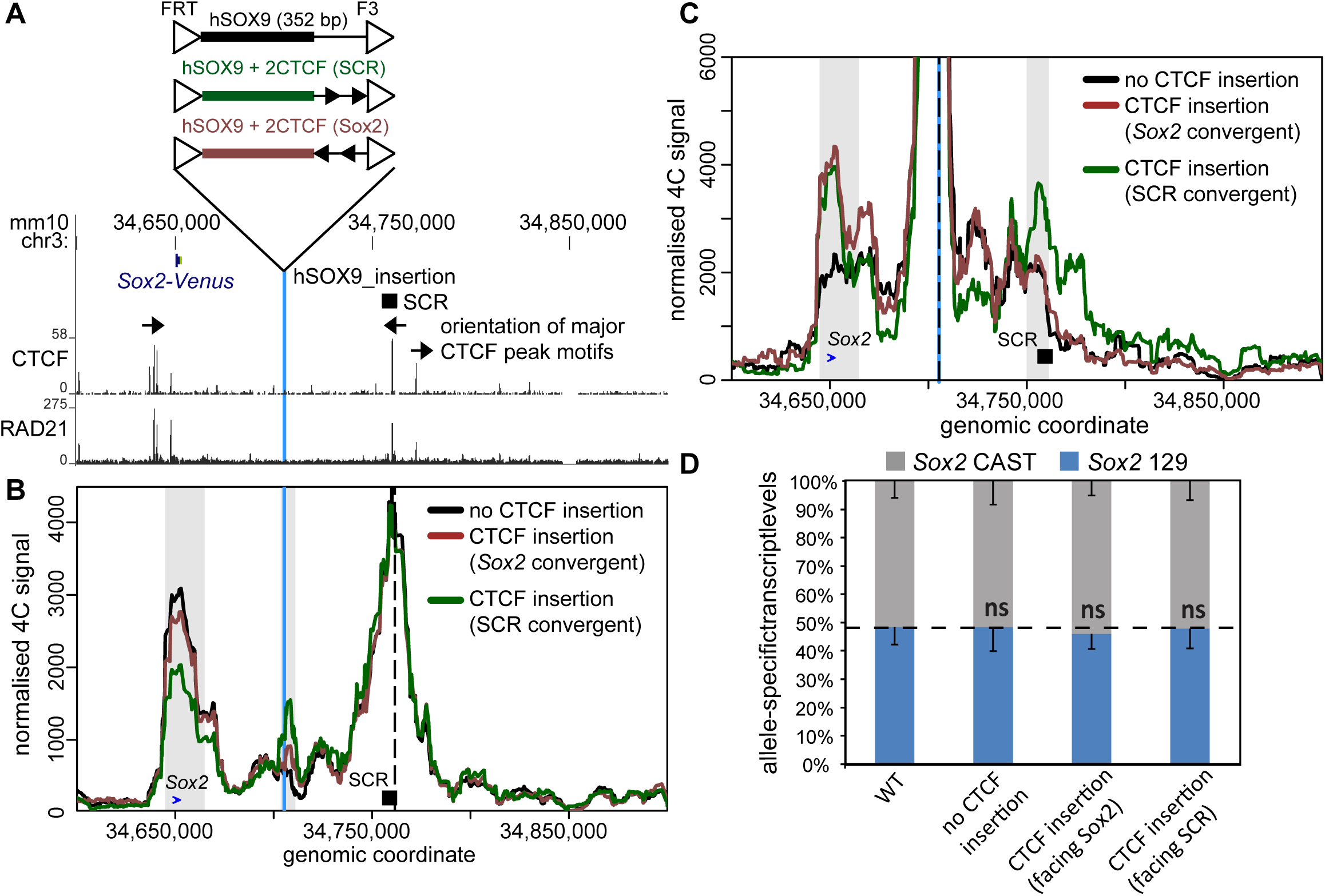
A CTCF motif insertion between the *Sox2* gene and the SCR disrupts the SCR­*Sox2* gene interaction, but not SCR-mediated enhancer activity. A) The region surrounding the *Sox2* gene is displayed on the UCSC Genome Browser (mm10). The *Sox2* control region (SCR) is shown along with the orientation of the CTCF motifs within CTCF bound regions. The vertical blue line represents the location into which the sequences shown above were integrated. CTCF and RAD21 ChIP-seq conducted in ESCs is shown below. In B-C, 4C data are shown for no CTCF integration (black), CTCF sites integrated in an orientation convergent with the SCR site (green), and CTCF sites integrated in an orientation convergent with the *Sox2* site (brown). The vertical blue line represents the location into which the sequences were integrated and the dashed line marks the location of the 4C bait at the SCR (B) or the integration site (C). Grey boxes indicate the bait-interacting regions where significant differences were identified surrounding the *Sox2* gene (blue arrow), SCR region (black box) or integration site (blue line). Compared to control cells lacking CTCF at the insertion site, a significant decrease in relative interaction frequency of the 4C bait region at the SCR with the *Sox2* gene was observed in the SCR convergent CTCF insertion (*P* = 0.002), but not the *Sox2* convergent CTCF insertion (*P* = 0.1). Compared to control cells, a significant increase in relative interaction frequency of the 4C bait region at the insertion with the *Sox2* gene site was observed in the SCR convergent CTCF insertion (*P* = 0.02), and the *Sox2* convergent CTCF insertion (*P* = 0.005). A significant increase in relative interaction frequency of the 4C bait region at the insertion with the SCR was observed in the SCR convergent CTCF insertion (*P* = 0.02), but not the *Sox2* convergent CTCF insertion (*P* = 0.6). D) Allele-specific primers detect *Sox2* musculus (129) or castaneus (CAST) mRNA by RT-qPCR from F1 ESC clones from the genotype indicated. Expression levels from either allele are shown relative to the total transcript levels. Error bars represent SD, n = 3. (ns = not significant; *P* > 0.05).

We engineered a short human sequence insertion (352 bp *Dpn*II-*Csp*6I fragment upstream of the *SOX9* gene) which could be used as a unique 4C bait. We introduced this insertion, with or without two copies of the core 19-bp CTCF motif within SRR109, into the landing site between *Sox2* and the SCR; the motifs were either both in convergent orientation with SRR109/SCR or both in convergent orientation with the CTCF site at the *Sox2* promoter (**Fig 6A**). With this setup, we were able to perform allele-specific 4C using either the SCR (**Fig 6B**) or the insertion site (**Fig 6C**) as the 4C bait. Insertion of the human bait without accompanying CTCF sites had no effect on endogenous *Sox2*-SCR interaction frequencies (**Fig S7C**). The insertion site did not interact with either the gene or the enhancer, although the observed abrupt loss of contact frequencies just distally of these elements was in line with their delimiting the *Sox2* TAD in ESCs (**Fig 5A**). As may be expected from the loop extrusion model, inclusion of CTCF sites facing *Sox2* created an ectopic interaction between the insertion site and the gene (*P* = 0.005), but not with the SCR, from either SCR or insertion site bait viewpoint (**Fig 6B, C**; **Table S1**). The endogenous *Sox2*-SCR interaction was unaffected (*P* = 0.1).

Inclusion of CTCF sites oriented towards the SCR created an ectopic interaction between the insertion site and the SCR (*P* = 0.02), which was clearly detected from both SCR and insertion site bait perspectives (**Fig 6B, C**). This profile was notably also accompanied by a 31% reduction in endogenous *Sox2*-SCR interaction frequencies, which is both significantly different from our control (*P* = 0.002) and quantitatively very similar to the interaction loss caused by deletion of the SCR. Surprisingly, the ectopic CTCF site also formed a strong and significant interaction with the *Sox2* promoter *(P* = 0.02), which would not have been expected from a loop extrusion model due to the incompatible motif orientations at these regions. Importantly, no allelic imbalance of *Sox2* expression was present in this or any of the other tested insertion-carrying line(s) (**Fig 6D**). These findings suggest that perturbation of chromatin architecture to the same extent as that caused by functionally significant enhancer deletions does not necessarily affect transcriptional output. Overall, this complementary approach reinforces the finding that the *Sox2*-SCR interaction appears very robust in ESCs, likely mediated by the contribution of many distributed elements, but that regulation of transcription and chromatin architecture is nonetheless highly decoupled.

## Discussion

We have performed extensive allele-specific engineering at the mouse *Sox2* locus to functionally dissect the contributions of different regulatory elements to transcriptional regulation and modulation of chromatin architecture in ESCs. In so doing, we have uncovered a striking decoupling of the two processes. On the one hand, the vast majority of distal *Sox2* transcriptional regulation can be ascribed to a small number of transcription factor-bound regions, whereas maintenance of the 3D architecture of the locus is very robust and seemingly distributed over large regions within the TAD. The allele-specific application in our study was critical to dissecting the *cis-*contributions of regulatory elements, since any small clonal variations were quantitatively controlled relative to the wild-type allele, and confounding *trans* effects of loss of SOX2 protein were avoided (Huang et al., 2021; Vos et al., 2021).

Based on its relatively large size and clustering of binding sites for tissue-specific transcription factors, the SCR fits the canonical description of a “super-enhancer” (Whyte et al., 2013). As such, the SCR is presumed to strongly activate target genes due to the combinatorial action of its composite transcription factor binding sites. However, the functional significance of “super-enhancers” compared to non-clustered enhancers is markedly debated, with previous genetic dissections uncovering context-dependent transcriptional effects that may be functionally redundant or additive, but not synergistic (Hay et al., 2016; Moorthy et al., 2017; Saravanan et al., 2020; discussed in Blobel et al., 2021). Although the 7.3 kb-long SCR contains four regions, each bound by more than six different transcription factors, we found that only two of these regions (SRR107 and SRR111) account for the vast majority of *Sox2* transcription regulation in *cis.* These two elements are each bound by eight or more transcription factors; however, their transcription-enhancing capacity can be greatly disrupted by removal of three motifs, a POU5F1:SOX2 composite motif in SRR107 and two KLF4 motifs in SRR111, together totaling 37 nucleotides. We recently showed a similar dependence on the POU5F1:SOX2 composite motif, as well as many other motifs for activity of the SRR107 region in vector-based reporter assays (Singh et al. 2021). In contrast, chromatin architecture at this locus is not dependent on only a few nucleotides, as only deletions of the entire SCR or even larger regions displayed significant disruptions to the interaction frequencies with *Sox2* or the TAD boundary downstream of the SCR. The two enhancers within the SCR seem to be at least partially redundant, since disruption of both SRR107 and SRR111 is required for a strong transcriptional defect. None of the multiple other transcription factor-bound regions within or outside the SCR appear to offer any functional redundancy in ESCs from a transcriptional standpoint. Since the SCR adopts an inactive chromatin state and genome architecture in later developmental stages (Ben Zouari et al., 2020; Bonev et al., 2017), these regions are also not expected to act as enhancers in differentiated cells. However, since naïve state ESCs are an *in vitro* approximation of pre-implantation epiblast cells (Avilion et al., 2003; Evans and Kaufman, 1981), these elements may regulate *Sox2* expression in different physiological contexts of pluripotent cells *in vivo*. Alternatively, but not mutually exclusively, the “surplus” landing sites for transcription factors may indirectly facilitate the initiation of 3D folding of the locus during development to mediate transcriptional activation by the major regulatory elements.

A current question in the field is to what extent any interplay between CTCF-stalled cohesin-mediated loop extrusion and protein-protein interactions between transcription factors bound at distributed genomic sites affects chromosome folding (Dowen et al., 2014; Schuijers et al., 2018; Kubo et al., 2021). In this study, we assessed this question at the *Sox2* locus. The SCR contains one site within SRR109 that is strongly bound by CTCF and whose motif is oriented towards the CTCF sites just upstream of the *Sox2* promoter, rendering this site an ideal candidate mediator of promoter-enhancer interactions via stalled loop extrusion. However, in line with a previous study (de Wit et al., 2015), SRR109 is completely dispensable for *Sox2* transcription and SCR-*Sox2* interaction. Additionally, we show that removal of this single CTCF-bound region or acute depletion of CTCF protein is not sufficient to disrupt the TAD boundary downstream of the SCR, although deletion of the entire SCR does cause loss of topological insulation. Importantly, insertion of the SRR109 CTCF motif into other genomic locations is sufficient to induce ectopic chromatin loops; therefore, the absence of a corresponding phenotype from the SRR109 deletion is not due to a lack of the strength of the motif within this *cis*-sequence. Instead, previous studies have shown a genomic context-dependent response of TAD borders to sequence deletions. Deletion of one or two CTCF sites at some loci, such as *Hoxa* and *Prdm14*, is sufficient for loss of TAD insulation and ectopic gene expression (Narendra et al., 2015; Vos et al., 2021), whereas other TAD borders, such as those at *Hoxd* and *Sox9/Kcnj2,* are highly resilient to large genomic deletions (Despang et al., 2019; Rodríguez-Carballo et al., 2017). At the *Sox2* locus, TAD border insulation is disrupted by the loss of the entire SCR, but not by the loss of its CTCF site, even when combined with the loss of both enhancers. We note that a parallel study prepared at the same time as our work also found the SCR CTCF site dispensable for both *Sox2* interaction and border function *in vivo* (Chakraborty et al., 2022). Interestingly, the loss of the *Sox2* promoter-proximal CTCF sites also had no effect on interaction frequencies with the SCR, but the TAD border upstream of the gene was perturbed. As well as further demonstrating the context-dependent nature of CTCF-mediated topological insulation, this apparent hierarchy of borders as “interaction attractors” could explain why an inserted intra-TAD CTCF site in either orientation ectopically interacted with the *Sox2* gene. The authors for this study concluded that strong enhancer-promoter interactions could “bypass” topological insulation instructed by CTCF, but the CTCF-independent role of the SCR in TAD border maintenance was not assessed. We went further to show that the interaction and domain organization at *Sox2* is completely independent of CTCF and transcriptional activation.

With the negligible role of CTCF in orchestrating chromatin topology at the *Sox2* locus, other mechanisms could mediate enhancer-promoter interactions and/or TAD organization, such as: cohesin recruitment at transcription factor-bound sites to initiate loop extrusion events (Liu et al., 2021; Vos et al., 2021); transcription factor and co-activator protein-protein interactions (Deng et al., 2012; Sabari et al., 2018); and ongoing transcription (Rowley et al., 2017; Hsieh et al., 2020). Strikingly, chromatin topology appears completely unaltered upon the compound deletion of SRR107 and SRR111, where transcription from the deleted allele is almost completely abolished. This result demonstrates a near-full uncoupling of transcriptional control and architecture maintenance by these two elements. Thus, ongoing transcription appears to have no role in *Sox2* chromatin topology, although a potential role for a low level of basal transcription cannot be ruled out. Instead, a role for transcription factor-bound sites themselves in mediating chromatin interactions is supported by a progressive, quantitative reduction of interaction as more transcription factor-bound regions are removed: no effect upon the deletion of SRR109 or SRR107 and SRR111; a weak, insignificant reduction upon removing all sequences between and including SRR107 and SRR111 (including SRR109); a significant reduction upon removal of the entire SCR; and weak further losses of interaction frequencies when also removing transcription factor binding sites at SRR85-SRR95 and/or an extension of the SCR deletion to a distal CTCF site. These data support the notion of a distribution of chromatin architecture maintenance over many genomic elements (presumably transcription factor-bound regions), within the *Sox2* TAD, each individually posing a weak effect but one that collectively builds up the domain. Such a result is consistent with previous findings that distal regulatory elements can co-associate in nuclear “hubs” (Allahyar et al., 2018; Espinola et al., 2021; Oudelaar et al., 2018; Palstra et al., 2003), often involving clusters of specific transcription factors (Li, J., et al., 2020; Papantonis et al., 2012; Schoenfelder et al., 2010). Furthermore, live imaging experiments have identified such a cluster around the *Sox2* transcription site, comprising co-associated SOX2, BRD4 and RNA polymerase II (Li, J., et al., 2020). We thus propose that the ESC-specific *Sox2* TAD is established as a consequence of such clustering, delimited by the *Sox2* promoter and the SCR, which exhibits the greatest transcription factor site density. Other loci, such as *Pou5f1*, may be similarly coordinated (Li, J., et al., 2020), whereas imaging experiments support a predominantly CTCF-mediated mechanism for other tissue-specific chromosome domains, such as the mouse *alpha-globin* locus (Brown et al., 2018).

The requirement for spatial proximity of promoters and enhancers for transcriptional control has recently been questioned by imaging experiments showing gene transcription in the absence of enhancer co-association (Alexander et al., 2019; Benabdallah et al., 2019). Whether the same observation applies to *Sox2* remains unclear. Consistent and large distances between *Sox2* and the SCR reported after imaging when large operator sequences are inserted near the elements (Alexander et al., 2019), are not in agreement with frequent close proximities measured by DNA FISH in wild-type fixed cells (Huang et al., 2021). Our own and others’ studies identify an ESC-specific proximity between *Sox2* and the SCR as demonstrated in population-averaged 3C-based studies (Bonev et al., 2017; de Wit et al., 2015; Zhou et al., 2014). However, this is not sufficient proof for a causative link between chromatin contact and transcriptional regulation. Notably, none of our own or parallel attempts to disrupt SCR-*Sox2* interactions via ectopic CTCF-mediated loops (**Figure 6**; Chakraborty et al., 2022; Huang et al., 2021) were able to completely abolish the endogenous promoter-enhancer interactions; therefore, the dependence of the interaction for transcription could not be assessed. Taken together, the seemingly conflicting reports from 3C-based and imaging studies suggest at least some form of spatial proximity between promoter and enhancer at some point in the transcription cycle. It has been proposed that hubs of transcription factors may generate unique nuclear microenvironments competent for transcription, perhaps within phase-separated condensates (Lim and Levine, 2021). Such environments would facilitate interactions between genes and distal genomic elements; although the increased local concentration of regulatory factors means that their juxtaposition would not be a strict prerequisite for transcriptional firing, but simply a co-association within the same hub. Recent studies are beginning to dissect the relative requirements of phase separation and conventional protein-protein dimerization events in building these hubs (Chong et al., 2018; Li, J., et al., 2020; Wei et al., 2020).

Despite the attractiveness of the transcription factor hub model, it should be noted that even upon removal of six regions within the *Sox2* TAD, each bound by multiple transcription factors and accounting for 49 separate binding events, the domain structure appears robust. More than half of the detected interaction frequency between *Sox2* and the 4C bait adjacent to the SCR remains after these deletions, leaving open the possibility that additional mechanisms may contribute to local chromatin topology. Previous studies have shown that abolition of the *Sox2*­SCR TAD upon ESC differentiation to neuronal precursors (Bonev et al., 2017) was associated with a complete loss of promoter-enhancer interactions within only a few days of *in vitro* differentiation (Ben Zouari et al., 2020). These data suggest that any such mechanisms are still tissue-specific and not “hard-wired” at this locus. Specifically, the TAD border upstream of the *Sox2* promoter, with four promoter-associated CTCF sites, is maintained, but all topological insulation at the SCR is lost. Notably, the seemingly dispensable CTCF binding event at SRR109 is likewise lost (Bonev et al., 2017), but this is also accompanied by a downregulation of the majority of pluripotency transcription factors that cluster at the SCR in ESCs (Dhaliwal et al., 2018). Combined with our own findings that local chromatin architecture is uncoupled from *Sox2* transcription and CTCF binding, the potential importance of spatial transcription factor clustering has not been fully established, and warrants further investigation.

Overall, our results reveal that specific elements supplying transcriptional enhancer function contribute to, but are not required for, maintained chromatin structure. Instead, we identify a distributed role of multiple regulatory elements in organizing chromatin structure, in stark contrast to the small number of elements necessary for transcriptional regulation. The dispensability of CTCF motifs (and protein) in the *Sox2* locus for both chromatin interaction and boundary maintenance highlights the shortcomings of focusing on CTCF over other transcription factor binding events that can similarly contribute to promoter-enhancer interactions and TAD boundary maintenance.

## Acknowledgments

We thank Barbara Panning and Marie-Christine Birling for sharing resources, and all members of the Sexton and Mitchell labs for critical reading of the manuscript. Sequencing was performed by the IGBMC GenomEast platform, a member of the France Génomique consortium (ANR-10-INBS-0009). This study was made possible because of the IGBMC flow cytometry and molecular biology platforms. Work in the Sexton lab was supported by funds from the European Research Council (ERC) under the European Union’s Horizon 2020 research and innovation program (Starting Grant 678624 - CHROMTOPOLOGY), the ATIP-Avenir program, and the grant ANR-10-LABX-0030-INRT, a French State fund managed by the Agence Nationale de la Recherche under the frame program Investissements d’Avenir ANR-10-IDEX-0002-02. NS is a fellow of the Fondation de Recherche Medicale. TS is supported by INSERM. Work in the Mitchell lab was supported by the Canadian Institutes of Health Research (FRN 153186), the National Institutes of Health (R01-HG010045-01), the Canada Foundation for Innovation, and the Ontario Ministry of Research and Innovation. VMS is supported by the NSERC Canadian Graduate Scholarship (CGS D).

## Author contributions

TT, NS, VMS, JAM and TS conceived and designed the study. TT, VMS, CJ and NNM created deleted ESC lines and analyzed gene expression. SM developed the *Sox2* allele tagging approach and maintained ESC lines. NS and SC performed 4C, with VMS generating some of the fixed chromatin. NS generated and characterized the ectopic CTCF ESC lines, with initial design input from SC. TS performed 4C analysis. All authors contributed to writing of the manuscript.

## Materials and Methods

### Cell culture

Mouse F1 ESCs (*M. musculus*^129^ × *M. castaneus*, female cells obtained from Barbara Panning) were cultured on 0.1% gelatin-coated plates in ES medium (DMEM containing 15% FBS, 0.1 mM MEM nonessential amino-acids, 1 mM sodium pyruvate, 2 mM GlutaMAX, 0.1 mM 2-mercaptoethanol, 1000 U/mL LIF, 3 µM CHIR99021 [GSK3β inhibitor; Biovision], and 1 µM PD0325901 [MEK inhibitor; Invitrogen]), which maintains ESCs in a pluripotent state in the absence of a feeder layer (Mlynarczyk-Evans et al., 2006; Ying et al., 2008).

### Cas9-mediated deletion

Cas9-mediated deletions were carried out as previously described (Moorthy and Mitchell, 2016; Zhou et al., 2014). Cas9 targeting guide RNAs (gRNAs) were selected flanking the desired region identified for deletion (**Table S2**). For select cases of allele-specific targeting, gRNA pairs were designed so that at least one gRNA overlapped a SNP to specifically target the *M. musculus*^129^ allele. On and off-target specificity of the gRNAs were calculated as described in (Doench et al. 2016; and Hsu et al., 2013) respectively to choose optimal guides. Guide RNA plasmids were assembled with gRNA sequences using the protocol described by Mali et al., (2013). Briefly, two partially complementary 61-bp oligos were annealed and extended using Phusion polymerase (New England Biolabs). The resulting 100-bp fragment was assembled into an *Afl*II-linearized gRNA empty vector (Addgene, ID#41824) using the Infusion assembly protocol (TaKaRa Bio). The sequence of the resulting guide gRNA plasmid was confirmed by sequencing with either T7 or SP6 primers.

F1 ESCs were transfected with 5 µg each of the 5′ gRNA(s), 3′ gRNA(s), and pCas9_GFP (Addgene, ID#44719) (Ding et al., 2013) or pCas9_D10A_GFP (Addgene, ID#44720) plasmids using the Neon Transfection System (Life Technologies). Forty-eight hours post-transfection, GFP-positive cells were isolated on a BD FACSAria. Ten thousand GFP positive cells were seeded on 10-cm gelatinized culture plates and grown for 5–6 days until large individual colonies formed. Colonies were picked and propagated for genotyping and gene expression analysis as previously described (Moorthy and Mitchell, 2016; Zhou et al., 2014). Genotyping of the deletions was performed by amplifying products internal to and surrounding the target deletion. All deleted clones identified from the initial screen were sequenced across the deletion; SNPs confirmed allele-specificity of the deletion (**Table S3**).

### Generation of insertion lines

A P2A-Venus reporter construct was inserted at *Sox2* on the musculus allele of F1 ESCs by homologous recombination after Cas9-mediated DNA break at the 3’ end of *Sox2*. A plasmid containing a P2A-Venus cassette and one containing Cas9-mCherry, a puromycin resistance gene and three gRNA cassettes were assembled by the IGBMC molecular biology platform (**Fig S6A**; vectors available on request). One gRNA targets a Cas9-mediated DNA break at the 3’ end of *Sox2*, and the other two target breaks flanking the P2A-Venus cassette on the plasmid to generate 8 bp microhomology to the *Sox2* 3’ site (**Fig S6A**). 5 µg of each plasmid was transfected into 1 million cells with lipofectamine 2000 and Venus-positive cells were isolated by FACS after five days. Single clones were isolated and incorporation of reporter into the musculus and/or castaneus allele was determined by allele-specific PCR. A musculus-incorporated heterozygous clone was further characterized by sequencing of PCR products. Expression of pluripotency markers, allelic *Sox2* expression and 4C chromatin interaction profiles were unperturbed by incorporation of the Venus reporter (**Fig S7**).

This reporter line was then used for musculus-specific insertion of an FRT/F3 cassette into a site between *Sox2* and the SCR. 1 kb homology arms were added to a plasmid containing a puromycin resistance-thymidine kinase positive-selection marker within an FRT/F3 cassette (kind gift from Marie-Christine Birling, Institut Clinique de la Souris) by restriction cloning (**Fig S6B)**. 5 µg of this plasmid was co-transfected with 5 µg Cas9-mCherry/sgRNA plasmid (generated by the IGBMC molecular biology platform; vectors available on request) into 1 million cells with lipofectamine 200. Only the musculus allele has a functional PAM at the gRNA target site due to a SNP. Cherry-positive cells were isolated by FACS after three days, and after one day of recovery, clones were selected with 3 µg/mL puromycin for one day, then 1 µg/mL puromycin until individual resistant colonies were formed. Clones were screened by PCR and confirmed by sequencing. We noted a slight reduction in musculus-specific *Sox2* transcription and SCR-*Sox2* interaction from this founder line, which was rescued on removal of the positive-negative selection marker (**Fig S7**).

The positive-negative selection marker was replaced with different inserts by FLP-mediated recombination. The donor vectors were constructed by restriction cloning of the initial FRT/F3 plasmid, annealed oligonucleotides (for CTCF motifs) and PCR products from human genomic DNA template (for SOX9 sequence). 5 µg donor vector was co-electroporated with 5 µg FLP-expressing plasmid (kind gift from Marie-Christine Birling, Institut Clinique de la Souris) with Neon, and after two days of recovery, recombinant clones were selected with 6 µM ganciclovir. Clones were screened by PCR and confirmed by sequencing. Flow cytometry quantitation revealed a slight but equal reduction in Venus reporter expression in all clones compared to the founder line (**Fig S7**), even though allelic balance of *Sox2* expression was unaltered (Fig 6D).

### RNA isolation and gene expression analysis by RT-qPCR

Total RNA was purified from single wells of >85% confluent six-well plates using the RNeasy Plus Mini Kit (Qiagen), and an additional DNase I step was used to remove genomic DNA. RNA was reverse-transcribed with random primers using the high-capacity cDNA synthesis kit (Thermo Fisher Scientific). *Sox2* gene expression was detected by allele-specific primers which specifically amplified either the musculus or castaneus allele as described in (Moorthy and Mitchell, 2016; Zhou et al., 2014). The standard curve method was used to calculate expression levels using F1 ESC genomic DNA to generate the standard curves. Levels of *Gapdh* or *Sdha* RNA were used to normalize expression values. Primer sequences are shown in **Table S5**.

### Allele-specific 4C-seq

Cells were fixed with 2% paraformaldehyde in 10% FBS in PBS for 10 min at 23°C. The fixation was quenched with cold glycine at a final concentration of 125 mM, then cells were washed with PBS and permeabilized on ice for 1 h with 10 mM Tris-HCl, pH 8, 100 mM NaCl, 0.1% NP-40, and protease inhibitors. Nuclei were resuspended in *Dpn*II restriction buffer at 10 million nuclei/mL concentration, and 5 million nuclei aliquots were further permeabilized by treatment for 1 h with 0.4% SDS at 37°C, then incubating for a further 1 h with 3.33% Triton X-100 at 37°C. Nuclei were digested overnight with 1500 U *Dpn*II at 37°C, then washed twice by centrifuging and resuspending in T4 DNA ligase buffer. *In situ* ligation was performed in 400 Μl T4 DNA ligase buffer with 20,000 U T4 DNA ligase overnight at 23°C. DNA was purified by reverse cross-linking with an overnight incubation at 65°C with proteinase K, followed by RNase A digestion, phenol/chloroform extraction, and isopropanol precipitation. The DNA was digested with 5 U/μg *Csp*6I at 37°C overnight, then re-purified by phenol/chloroform extraction and isopropanol precipitation. The DNA was then circularized by ligation with 200 U/μg T4 DNA ligase under dilute conditions (3 ng/μL DNA), and purified by phenol/chloroform extraction and isopropanol precipitation. For allele-specific 4C from the SCR-proximal bait, samples of the DNA were digested with *Bve*I or *Alw*26I, cutting specifically at the region between the *Dpn*II site and the 4C reading primer annealing site on the 129 or castaneus allele, respectively. For 129-specific 4C from the SCR bait, the material was digested with *Ava*III, cutting specifically at the region between the *Csp*6I site and the 4C non-reading primer annealing site on the castaneus allele. There were no SNPs allowing castaneus-specific 4C from this bait. No digestion was required for allele-specific 4C from the human insertion sequence, since this was only present on the 129 allele. 100 ng aliquots of treated DNA were then used as template for PCR with bait-specific primers containing Illumina adapter termini (primer sequences in **Table S6**). PCR reactions were pooled, primers removed by washing with 1.8× AMPure XP beads, then quantified on a Bioanalyzer (Agilent) before sequencing with a HiSeq 4000 (Illumina).

### 4C-seq analysis

Sequencing read fastq files were demultiplexed with sabre (https://github.com/najoshi/sabre) and aligned to the mm10 genome with Bowtie (Langmead et al., 2009), and intrachromosomal reads were assigned to *Dpn*II fragments by utility tools coming with the 4See package (Ben Zouari et al., 2020). 4See was also used to visualize the 4C profiles. Interactions were called for each replicate with peakC (Geeven et al., 2018) with window size set to 21 fragments, and were then filtered to only include the regions called as interacting across all wild-type replicates. We note that the “minimal” region spanning *Sox2* was called as interacting with the proximal SCR bait for all cell lines (and virtually all replicates) tested in this study (**Table S1**). For statistical comparison of specific interactions, the 4C read counts within 1 Mb of the bait for all replicates and conditions (from the same bait) were quantile normalized using the limma package (Ritchie et al., 2015). The means of summed normalized 4C scores over tested interacting regions were taken as “interaction scores”, and were compared across conditions by two-tailed t-tests. For the SCR-*Sox2* and insertion site-*Sox2* interactions, we used the minimal region spanning *Sox2* for all wild-type replicates with the near-SCR bait (mm10; chr3: 34,644,922-34,664,967). For the insertion site-SCR interaction, we used the minimal region spanning the SCR called as interacting in all replicates of the [CTCF insertion (SCR convergent)] line (mm10; chr3: 34,749,652-34,760,919). For quantifying inter-TAD interactions from the near-SCR bait, we used the 325 kb region (mm10; chr3: 34,800,000-35,105,000) which contains nearly the entire downstream TAD. Since 4C-seq signal at the 5’ end of the downstream TAD is artificially inflated in various deletion lines because the genomic separation has been shortened by the deletion, the selected region starts at a conservatively chosen place 3’ to 4C-seq local minima (i.e. when the contact decay with genomic separation has equilibrated for all the cell lines). As a control for the upstream TAD, the same size region was used (mm10; chr3: 34,315,000-34,640,000).

### Hi-C re-analysis

Published Hi-C data (Bonev et al., 2017; Nora et al., 2017) were downloaded from GEO (GSM2533818-2533821 for ESC *Dpn*II; GSM2533822-2533825 for NPC *Dpn*II; GSM2644945-2644946 for ESC control *Hind*III; GSM2644949-2644950 for ESC CTCF degron *Hind*III) and re-analysed using the FAN-C toolbox (Kruse et al., 2020), entailing read mapping, filtering out technical artefacts, mapping to restriction fragment space, binning, matrix normalisation and ratio-based comparison, and visualisation.

### ChIP-seq visualization

Published ESC ChIP-seq datasets (Arruda et al., 2020; Chen et al., 2008; Creyghton et al., 2010; Kagey et al., 2010; Wamstad et al., 2012) were downloaded from GEO (GSM4280484 for RAD21; GSM288351 for CTCF; GSM1163096 for H3K27ac; GSM594600 for EP300; GSM560348 for MED1) and visualized with the UCSC Genome Browser.

### Data availability

All 4C-seq data from this study have been deposited on GEO with the accession GSE195906.

