## Supplemental Figures and Tables for "Transcriptional regulation and chromatin architecture maintenance are decoupled modular functions at the *Sox2* locus"

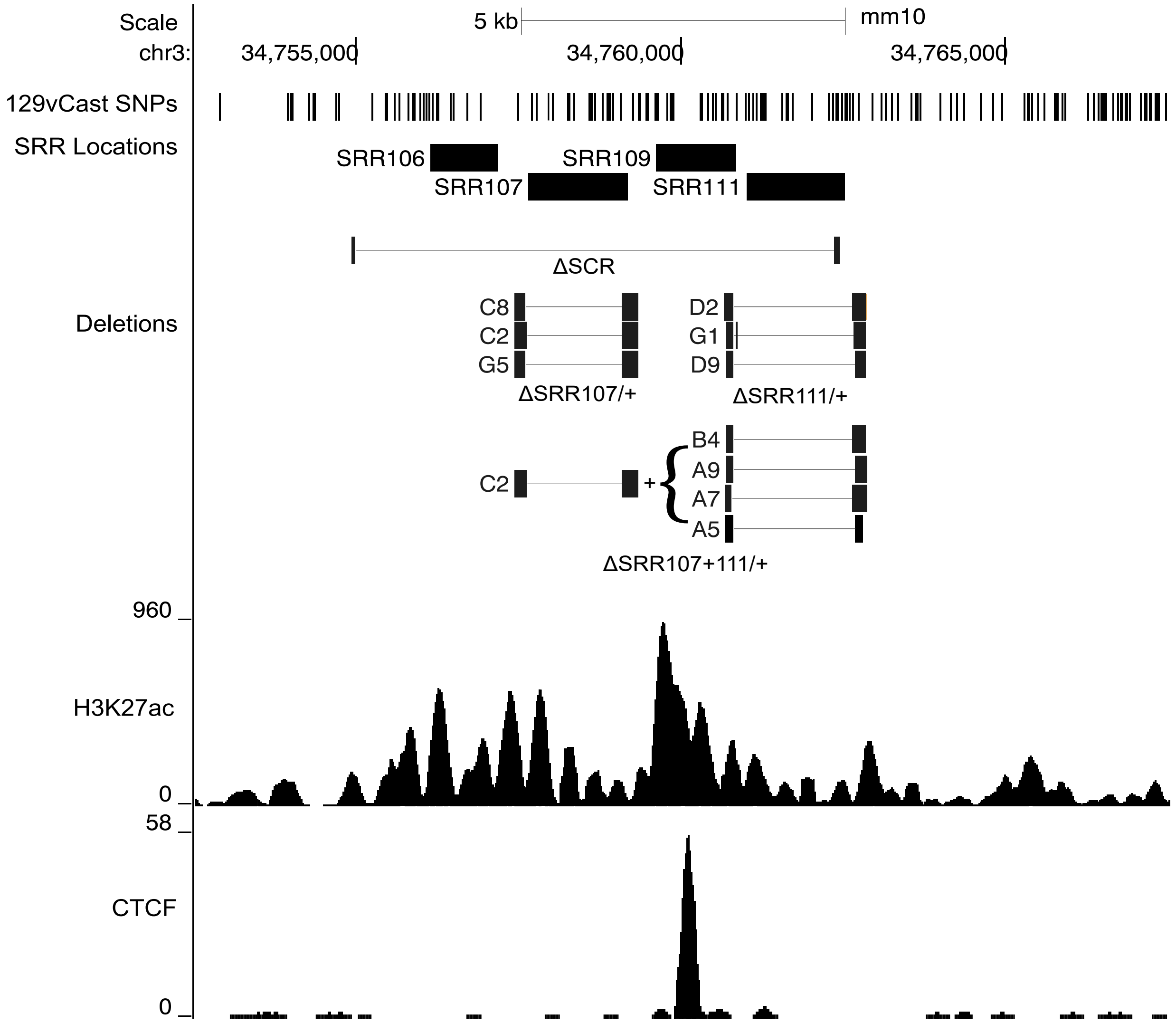

Scale
5 kb
mm10
chr3:
34,755,000
34,760,000
34,765,000
129vCast SNPs
SRR Locations
SRR106
SRR107
SRR109
SRR111
ΔSCR
Deletions
C8
C2
G5
ΔSRR107/+
D2
G1
D9
ΔSRR111/+
C2
+
B4
A9
A7
A5
ΔSRR107+111/+
960
H3K27ac
0
58
CTCF
0

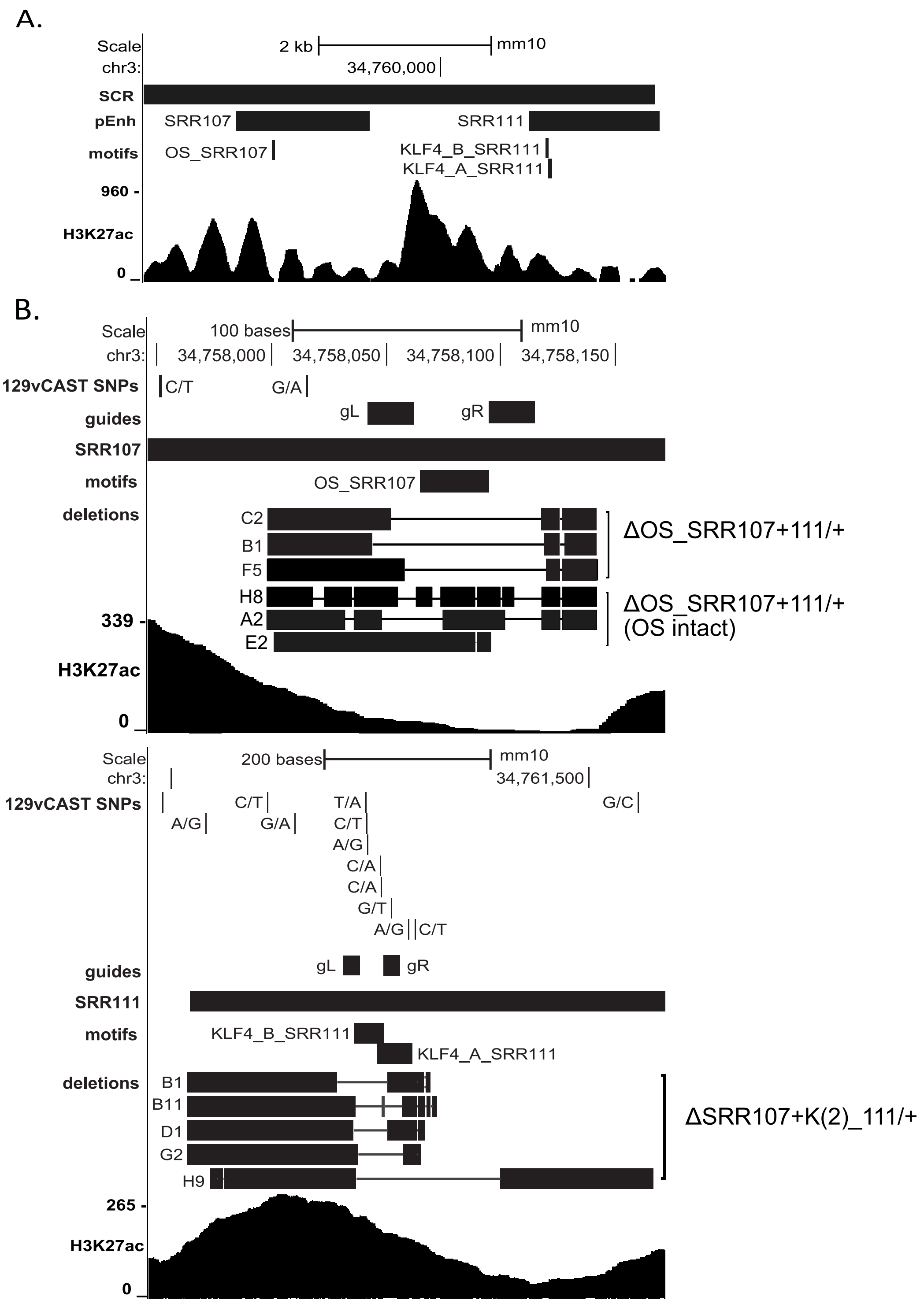
A.
Scale
2 kb
mm10
chr3:
34,760,000
SCR
pEnh
SRR107
SRR111
motifs
OS_SRR107
KLF4_B_SRR111
KLF4_A_SRR111
960 -
H3K27ac
0
B.
Scale
100 bases
mm10
chr3:
34,758,000
34,758,050
34,758,100
34,758,150
129vCAST SNPs
C/T
G/A
guides
gL
gR
SRR107
motifs
OS_SRR107
deletions
C2
B1
F5
H8
A2
E2
ΔOS_SRR107+111/+
ΔOS_SRR107+111/+
(OS intact)
339 -
H3K27ac
0
Scale
200 bases
mm10
chr3:
34,761,500
129vCAST SNPs
C/T
T/A
G/C
A/G
G/A
C/T
A/G
C/A
C/A
G/T
A/G
C/T
guides
gL
gR
SRR111
motifs
KLF4_B_SRR111
KLF4_A_SRR111
deletions
B1
B11
D1
G2
H9
ΔSRR107+K(2)_111/+
265 -
H3K27ac
0

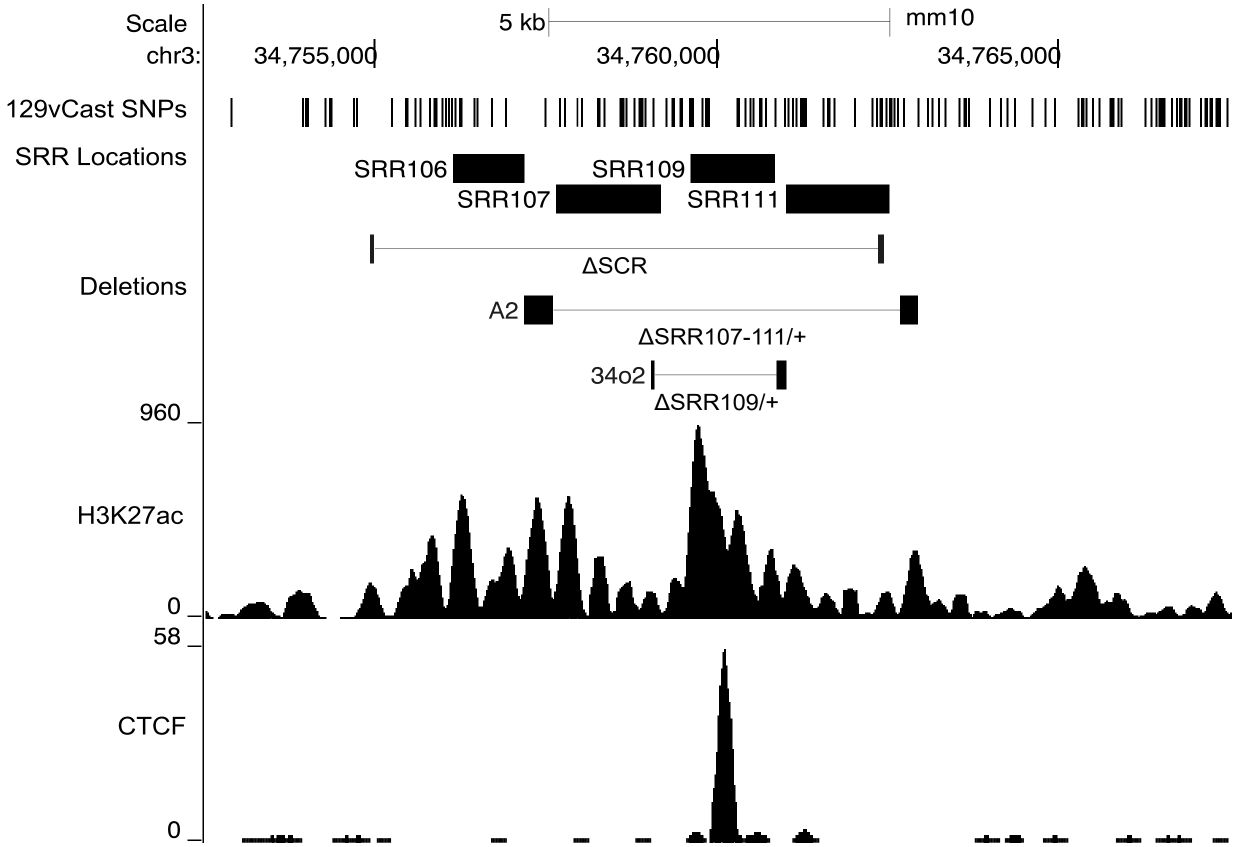
Scale
5 kb
mm10
chr3:
34,755,000
34,760,000
34,765,000
129vCast SNPs
SRR Locations
SRR106
SRR107
SRR109
SRR111
Deletions
ΔSCR
A2
ΔSRR107-111/+
34o2
ΔSRR109/+
960
H3K27ac
0
58
CTCF
0

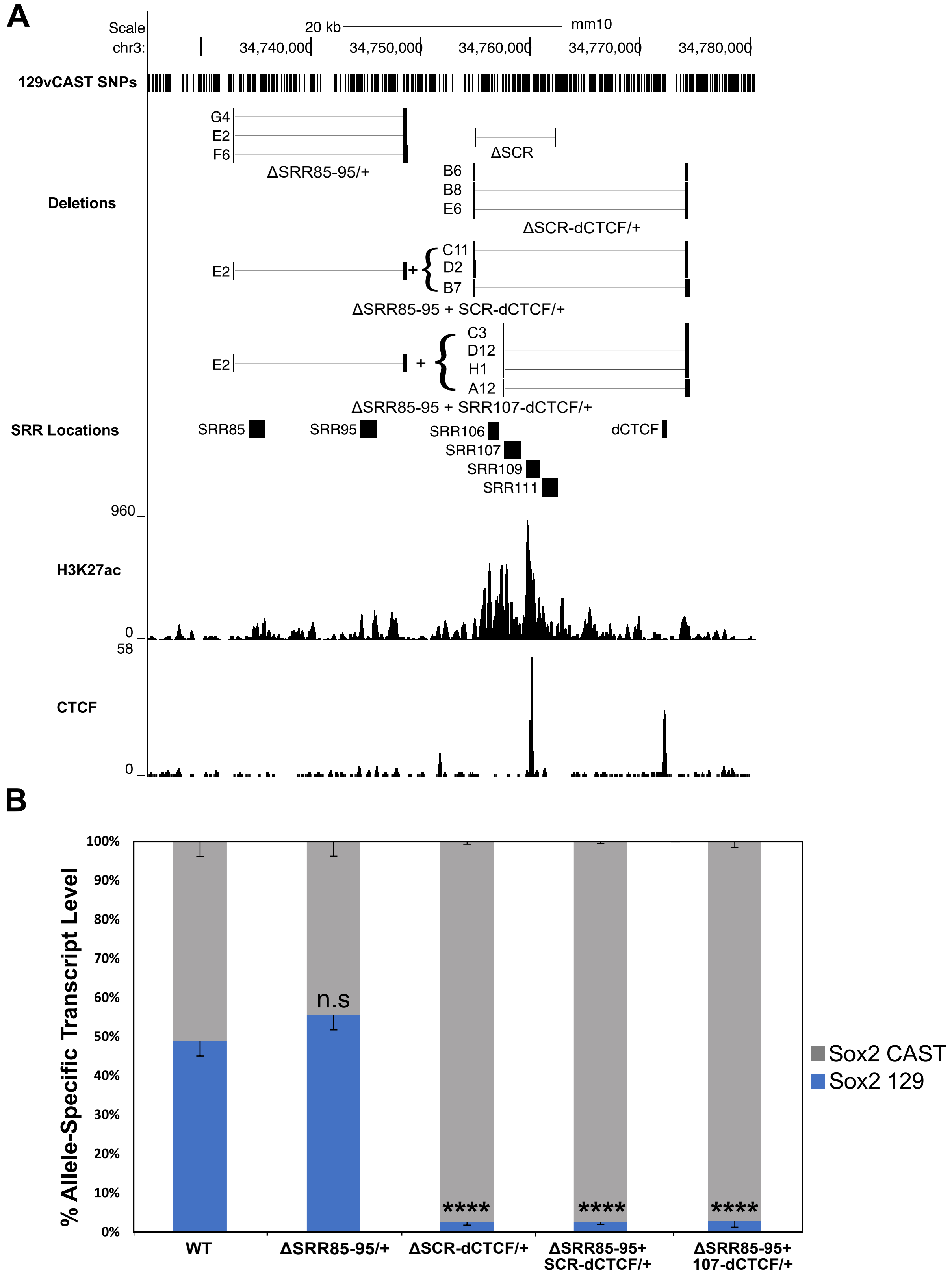

A
Scale
20 kb
mm10
chr3:
34,740,000
34,750,000
34,760,000
34,770,000
34,780,000
129vCAST SNPs
Deletions
G4
E2
F6
ΔSRR85-95/+
ΔSCR
B6
B8
E6
ΔSCR-dCTCF/+
E2
C11
D2
B7
ΔSRR85-95 + SCR-dCTCF/+
E2
C3
D12
H1
A12
ΔSRR85-95 + SRR107-dCTCF/+
SRR Locations
SRR85
SRR95
SRR106
SRR107
SRR109
SRR111
dCTCF
960
H3K27ac
0
58
CTCF
0
B
% Allele-Specific Transcript Level
100%
90%
80%
70%
60%
50%
40%
30%
20%
10%
0%
n.s
****
****
****
WT
ΔSRR85-95/+
ΔSCR-dCTCF/+
ΔSRR85-95+ SCR-dCTCF/+
ΔSRR85-95+ 107-dCTCF/+
Sox2 CAST
Sox2 129

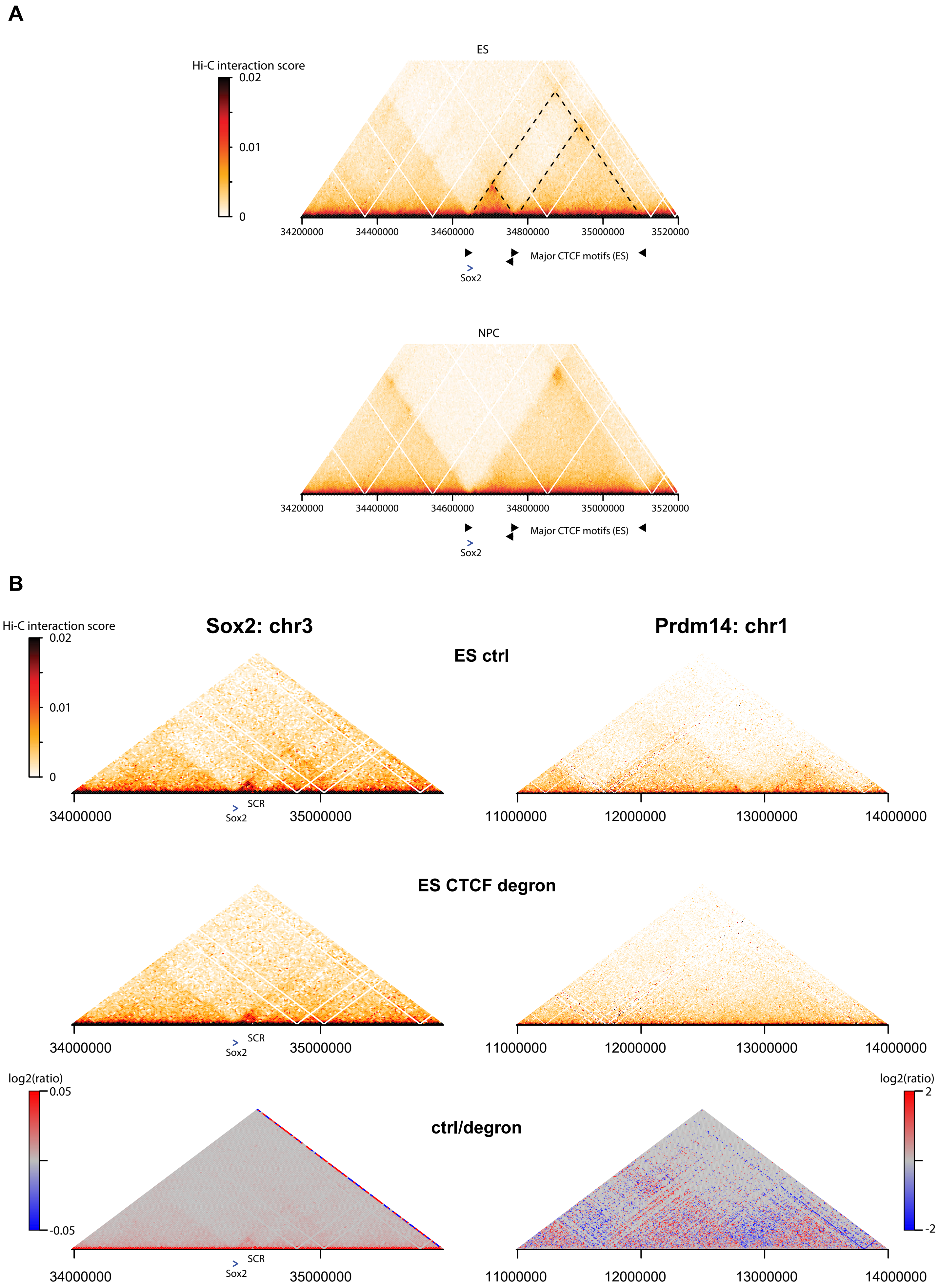
A
ES
Hi-C interaction score
0.02
0.01
0
34200000
34400000
34600000
34800000
35000000
3520000
Major CTCF motifs (ES)
Sox2
NPC
34200000
34400000
34600000
34800000
35000000
3520000
Major CTCF motifs (ES)
Sox2
B
Hi-C interaction score
0.02
0.01
0
Sox2: chr3
Prdm14: chr1
ES ctrl
SCR
Sox2
34000000
35000000
11000000
12000000
13000000
14000000
ES CTCF degron
SCR
Sox2
34000000
35000000
11000000
12000000
13000000
14000000
log2(ratio)
0.05
-0.05
ctrl/degron
log2(ratio)
2
-2
SCR
Sox2
34000000
35000000
11000000
12000000
13000000
14000000

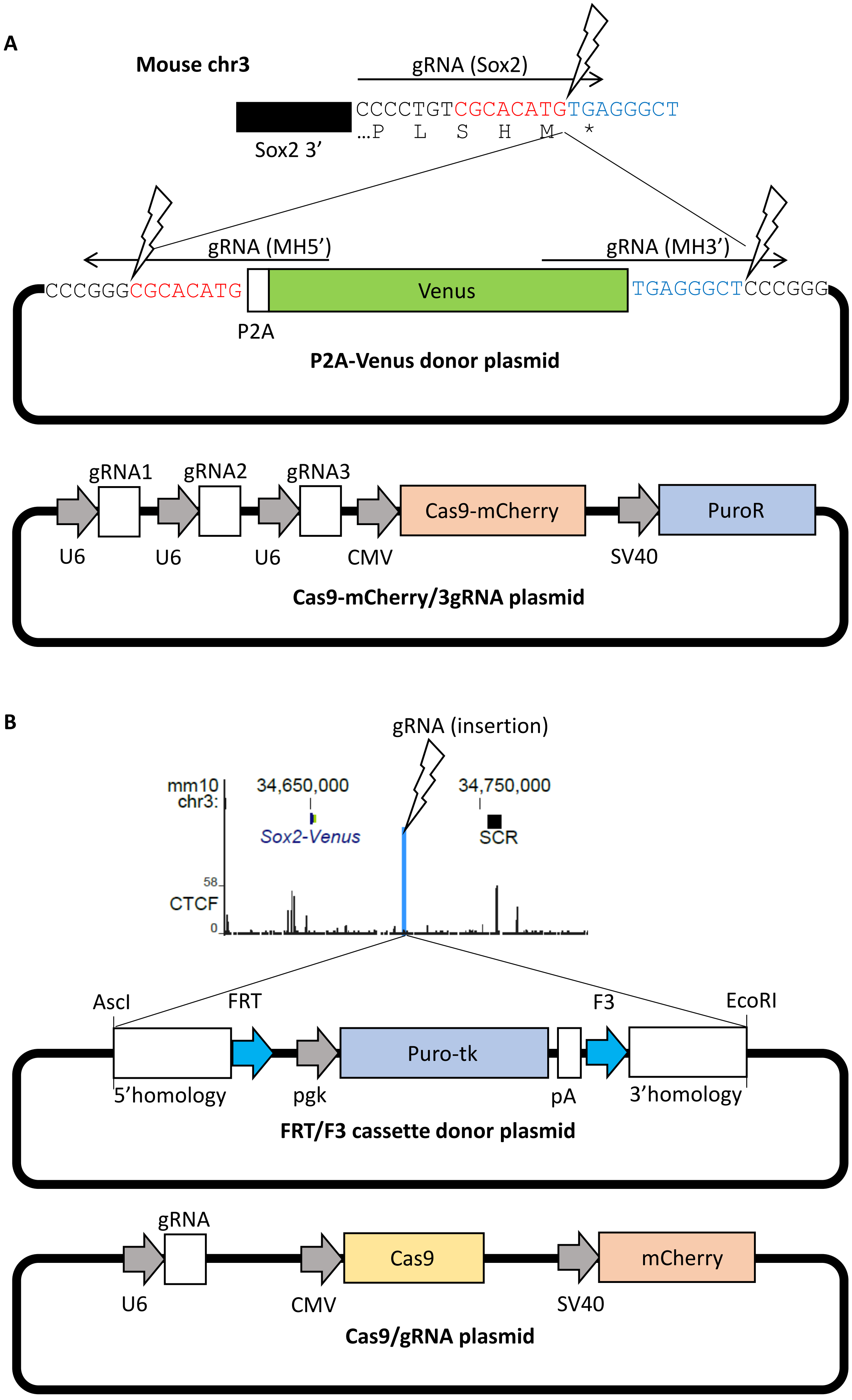
A
Mouse chr3
gRNA (Sox2)
CCCCTGTCGCACATGTGAGGGCT
...P L S H M *
Sox2 3'
gRNA (MH5')
gRNA (MH3')
CCCGGGCGCACATG
Venus
TGAGGGCTCCCGGG
P2A
P2A-Venus donor plasmid
gRNA1
gRNA2
gRNA3
Cas9-mCherry
PuroR
U6
U6
U6
CMV
SV40
Cas9-mCherry/3gRNA plasmid
B
gRNA (insertion)
mm10 chr3:
34,650,000
34,750,000
Sox2-Venus
SCR
58
CTCF
0
AscI
FRT
F3
EcoRI
Puro-tk
5'homology
pgk
pA
3'homology
FRT/F3 cassette donor plasmid
gRNA
Cas9
mCherry
U6
CMV
SV40
Cas9/gRNA plasmid

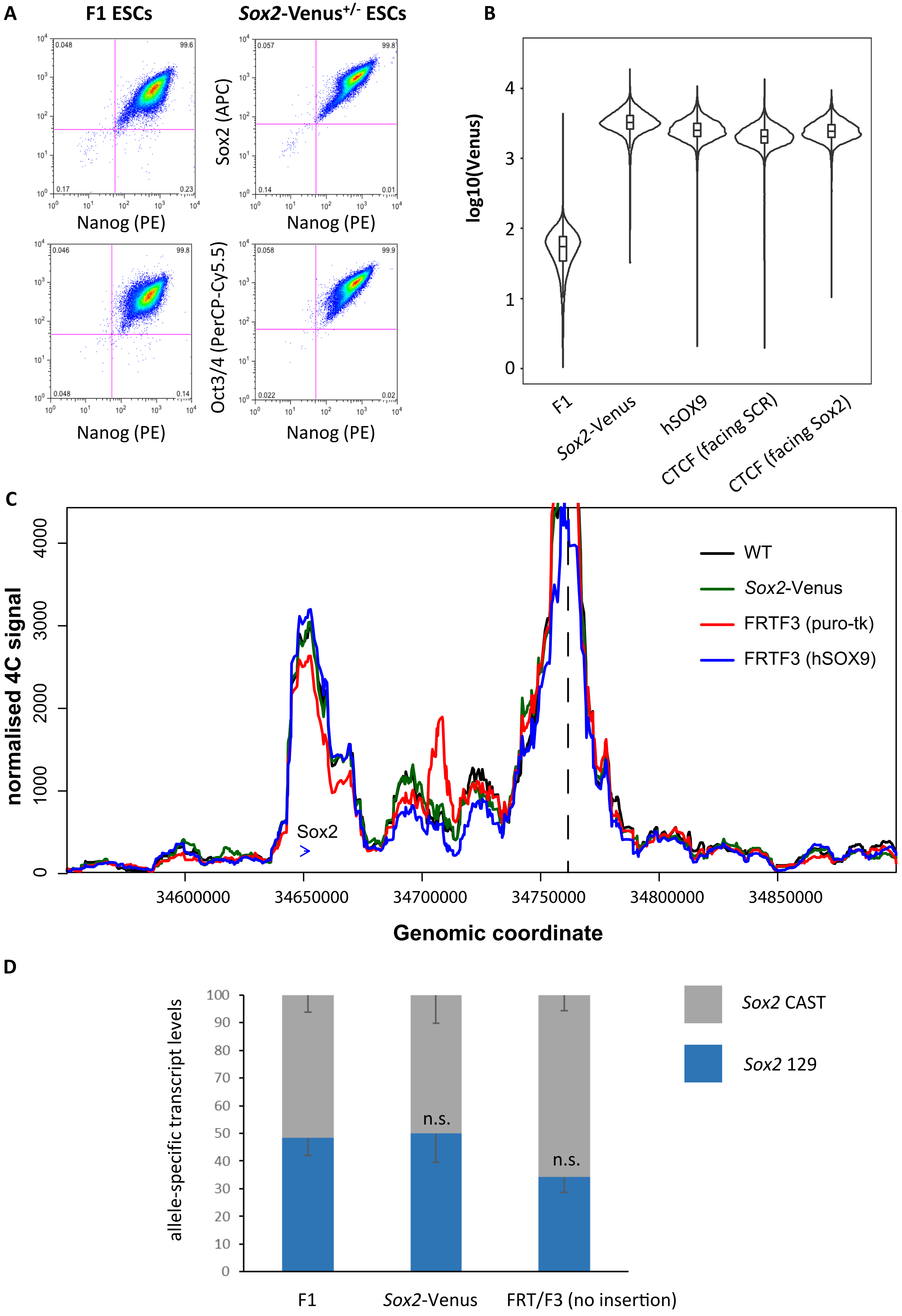
A
F1 ESCs
Sox2-Venus+/- ESCs
Sox2 (APC)
Oct3/4 (PerCP-Cy5.5)
Nanog (PE)
B
log10(Venus)
F1
Sox2-Venus
hSOX9
CTCF (facing SCR)
CTCF (facing Sox2)
C
normalised 4C signal
Genomic coordinate
Sox2
WT
Sox2-Venus
FRTF3 (puro-tk)
FRTF3 (hSOX9)
D
allele-specific transcript levels
Sox2 CAST
Sox2 129
n.s.
n.s.
F1
Sox2-Venus
FRT/F3 (no insertion)

### SUPPLEMENTARY FIGURE LEGENDS

**Supplementary Figure S1. Details on the generation of sub-SCR deleted cell lines targeting highly transcription factor-bound regions** Schematic representation of SCR region along with deletion boundaries displayed on the University of California Santa Cruz (UCSC) Genome Browser (mm10). Schematics explained from top to bottom: Genome coordinates, positions of *Mus castaneus* SNPs, positions of the SCR and SRR sub regions and the sequenced clones harbouring the specified deletion, ChIP-seq of H3K27ac and CTCF in wild-type mouse embryonic stem cells. For compound deletions ΔSRR107+111/+ clones were created in the background of ΔSRR107/+ C2. SCR deletion is shown as a comparison to the listed clones.

**Supplementary Figure S2. Details on the generation of sub-SCR deletions targeting specific transcription factor motifs.** A) Schematic representation of high-scoring predictive transcription factor motifs located in SRR107 or SRR111 of the SCR. SRR107 contains a high-scoring Oct:Sox motif (OS_SRR107) while SRR111 contains two high-scoring Klf4 motifs (KLF4_A_SRR111 and KLF4_B_SRR111, respectively) displayed on the UCSC Genome Browser (mm10). B) Resulting 129-specific deletions of targeted motif deletions are shown. Schematics explained from top to bottom: Genome coordinates, positions of *Mus castaneus* SNPs, positions of the SRR sub region, transcription factor motifs, the sequenced clones harbouring the specified deletion, ChIP-seq of H3K27ac in wild-type mouse embryonic stem cells.

**Supplementary Figure S3. Details on the generation of sub-SCR deleted cell lines targeting the central CTCF bound region and surrounding transcription factor-bound sites.** Schematic representation of the deletion encompassing both enhancer regions (SRR107 and SRR111) and the CTCF bound peak at SRR109 as well as the SRR109 region alone within the SCR displayed on the UCSC Genome Browser (mm10). Schematics explained from top to bottom: Genome coordinates, positions of *Mus castaneus* SNPs, positions of the SCR and SRR sub regions, the sequenced clones harbouring the specified deletion, ChIP-seq of H3K27ac and CTCF in wild-type mouse embryonic stem cells. SCR deletion is shown as a comparison to the listed clones.

**Supplementary Figure S4. Details on the generation of deletion lines including targets surrounding the SCR.** A) Schematic representation of the regions targeted for larger 129-specific deletions removing the SCR alongside a downstream CTCF bound region displayed on the UCSC Genome Browser (mm10). Schematics explained from top to bottom: Genome coordinates, positions of *Mus castaneus* SNPs, positions of the SCR and SRR sub regions, the sequenced clones harbouring the specified deletion, ChIP-seq of H3K27ac and CTCF in wild-type mouse embryonic stem cells. For compound deletions, clone Δ85-85/+ E2 was used to create ΔSRR85-95+SCR-dCTCF/+ and ΔSRR85-95+107-dCTCF/+. SCR deletion is shown as a comparison to the listed clones. B) *Sox2* expression in wild type F1 cells (WT) compared to clones carrying the indicated deletion on the 129 allele. Allele-specific primers detect musculus (129) or castaneus (CAST) RNA in RT-qPCR. Expression levels are normalized to transcript levels from GAPDH. Error bars represent the SD. Significant differences between wild-type cells and clones denoted by (*), with adjusted p-value <0.0001 (****), <0.001 (***), <0.01 (**), <0.05 (*) or non-significant (n.s.).

**Supplementary Figure S5. Chromatin architecture around the *Sox2* locus is ESC-specific and CTCF-independent.** A) Hi-C maps around the *Sox2* locus for ESCs (top) and neuronal precursor cells (NPC, bottom), generated from data from Bonev et al. (2017). Positions and orientations of the *Sox2* gene and major CTCF-bound motifs in ESCs are denoted under the maps. Dotted lines indicate ESC TADs. These two TADs fuse into one larger TAD in NPCs. B) Hi-C maps around the *Sox2* locus (left) and a control region around the *Prdm14* locus (right) in control ESCs (top) and ESCs after acute depletion of CTCF using an engineered auxin-inducible degron (middle); generated from data from Nora et al. (2017). Whereas clear loss of TADs on CTCF depletion is observed in the control region, *Sox2* architecture appears largely unchanged. Bottom: Quantitative comparison of the two experimental conditions at these regions is shown as a heatmap of log2(ctrl/degron) interaction scores. In the control region, CTCF depletion causes a ~4-fold relative increase in inter-TAD contacts and a similar decrease in intra-TAD contacts. At *Sox2,* contact changes are negligible (<1.04-fold).

**Supplementary Figure S6. Construction of *Sox2* insertion lines.** A) Plasmids constructed to generate the Venus tag at the 3' end of *Sox2*. F1 ESCs are transfected with a plasmid containing the P2A-Venus cassette (middle) and a plasmid (bottom) containing Cas9-mCherry, a puromycin resistance marker and expression constructs for three gRNAs: *Sox2* gRNA, which targets CRISPR/Cas9 to the 3' of the *Sox2* coding sequence (top), and MH5' and MH3' gRNAs, which target CRISPR/Cas9 to the 5' and 3' ends, respectively of the P2A-Venus cassette to generate 8 bp microhomology arms. B) Insertion of the recombinase-mediate cassette exchange construct. The *Sox2*-Venus$^{+/-}$ line is transfected with two plasmids: one (middle) containing an FRT-puro-tk-F3 cassette for positive-negative selection, flanked by homology arms for the *Sox2* intervening sequence, and one (bottom) containing Cas9, mCherry and a construct for expression of one gRNA, which targets CRISPR/Cas9 to a musculus site located between *Sox2* and the SCR (top).

**Supplementary Figure S7. Characterisation of the *Sox2* insertion lines.** A) Flow cytometry profiles of F1 (left) and *Sox2*-Venus$^{+/-}$ (right) ESCs after staining with labelled antibodies to Sox2, Nanog and Oct3/4, showing that stemness is unaffected on insertion of the Venus reporter tag. B) Flow cytometry quantitation of Venus fluorescence in the ESC lines with different insertions. Venus reporter is highly and equivalently expressed from lines where the hSOX9 tag, with or without CTCF sites, is inserted between *Sox2* and the SCR, which is slightly lower than the founder *Sox2*-Venus$^{+/-}$ line. C) *Musculus*-specific 4C profiles using the SCR as bait (dashed line) for F1 (black), *Sox2*-Venus$^{+/-}$ (green), FRT/F3/positive-negative selection marker (red), and FRT/F3/hSOX9 (blue) lines. The *Sox2*-SCR interaction is largely maintained, with a slight decrease in the presence of the positive-negative selection marker. D) Allele-specific qPCR quantitation of *Sox2* expression, relative to SHDA, for the musculus (129; blue) and castaneus (CAST; grey) alleles in F1, *Sox2*-Venus$^{+/-}$, and FRT/F3/positive-negative selection marker lines. Musculus *Sox2* transcription is weakly reduced in the presence of the selective marker. Error bars show SD (n = 2) (n.s. = non-significant).

### SUPPLEMENTARY TABLES

**Supplementary Table S1: Called interactions for each 4C-seq replicate experiment performed in this study.** Provided as a separate Excel file.

**Supplementary Table S2: Guide RNA sequences for CRISPR/Cas9 mediated deletions.**

ΔSCR guide RNAs are shown for clarity but were originally designed in Zhou *et al.,* 2014

| Target Region | Left Guide Sequence | Right Guide Sequence | Use with Cas9 or Cas9-D10A (nickase) |
|---|---|---|---|
| ΔSCR | TAGCATACGTCACGCCGGAA | ACTGTTCTCGAACACTCTGT | Cas9 |
| ΔSRR107-111 | GACAAAAACATGTACGTTGGG | GGCCAAGGTTGAGCTCTAGT | Cas9 |
| ΔSRR107 | GACAAAAACATGTACGTTGGG | CATTCCTTGGCCAGATGCTA | Cas9 |
| ΔSRR111 | CTTAAATTTTATTTTGTGCT | CTTGCTGAAGAGAACTAACC | Cas9-D10A |
| | TGGTCCCAGCATGTGCATA | GGCCAAGGTTGAGCTCTAGT | |
| ΔSRR109 | CATTCCTTGGCCAGATGCTA | GAGTTGAAAAGATGGCTCAG | Cas9-D10A |

| | | | |
|---|---|---|---|
| | ACATTGAACTAAGATCA TTT | CTTAAATTTTATTTTGTG CT | |
| ΔOct:Sox_1 07 | TAGTCCCAGGACTCTGCT AA | GGTGGGTAGTTAGCATA ATG | Cas9-D10A |
| ΔKlf4(x2)_ 111 | AGAAGATGAGATGAAAG GCA | TTGAAGGCAGCCTTCCGG TA | Cas9-D10A |
| ΔSRR85-95 | AACTTAGTGGACCATAC CCA | CAGTATGACACGCAGTG GCG | Cas9 |
| ΔSCR- dCTCF | TAGCATACGTCACGCCG GAA | GCTGCAAAGGCTCCCGTT CG | Cas9 |
| ΔSRR107- dCTCF | GAAGACAAAAACATGTA CGT | GCTGCAAAGGCTCCCGTT CG | Cas9 |

**Supplementary Table S3: Sequences across CRISPR/Cas9 mediated deletions.**

Removed region is denoted with a vertical line where CRISPR/Cas9 double-strand break or nick sites have been joined together. ΔSCR clones were created in Zhou *et al.,* 2014. All sequences are from the 129 allele except for specified ΔSCR clones. Targeted regions are also listed by their coordinates from the UCSC genome browser build mm10 based on gRNA locations.

| **Deletion Target (mm10)** | **Clone** | **Included in 4C Analysis (Y/N)** | **Deletion Sequence** |
|---|---|---|---|
| ΔSCR/ΔSCR(Cast) chr3: 34754958-34762355 | 1 | Y | AACTATAATTTCCTGTACAGTCTTTCTTTAGACAGGGCTT CTGCTGCCTCTGAGTGGAAGATTGCTGGGATCATATGCCA TCATATACATACCTGCACATATATGTGGGCACTTGCTGTA GGCAGAGGCCAGAAGAGGACAACACATCCCCTGGCTAGA GTTACAGTCAGCTCTGGAAAAAGCAGTAAGTGCTCTAAC CACTGAGCTACCGTTCCG\|TAGTCAGGGATGCACAGAGAA ACCGTCTCAAAAGACAAATACAGTAACCAAGACCAAAAC CAACAACCAAACCAAACCAAACACAAACCCCAAAACAA AAGGCTAAGCTATTCC |
| ΔSCR/ΔSCR (129) chr3: 34754958-34762355 | 1 | Y | CCCCTGGAAAAGCAGTAAGTGCTCTAACCACTGAGCTAC CGTTC\|GAGTGTTCGAGAACAGTCAGGGATGCACAGAGAA ACCGTCTCAAAAGACAAATACAATAACCAAGACCAAA |
| ΔSCR/+ (129) chr3: 34754958-34762355 | 15 | Y | TCCCCTGGAAAAGCAGTAAGTGCTCTAACCACTGAGCTA CCGTTC\|GAGTGTTCGAGAACAGTCAGGGATGCACAGAGA |

| | | | |
|---|---|---|---|
| | | | AACCGTCTCAAAAGACAAATACAATAACCAAGACCACCA<br>AC |
| +/ΔSCR (Cast)<br>chr3: 34754958-<br>34762355 | 11 | Y | ATTTAGAATTTTTAAAGAATTTTTATTTTATTTTTAATTTA<br>TTTTTTACACTTCATATTCCACTCCCTAACCCCCCATCATT<br>TAGAATTTTAATGGATTTCCTTCTATATATTTTTATTAATT<br>TTTTCTGTTTTAATTTTTTCTTTTTAAATTTTTCTTGTTTAA<br>TCTTTCTTAAATTCAATTTAAAAATTTTTAACATAAATAA<br>CTAATTTGTGTTAGTTATATGTAAACAGCGGCTTTCCCTG<br>TGTCATTTTCAGGCGTTTGTGTAGAGCATT\|CAAAAGAGG<br>GGGGGAAATGTAGATAATAATAAGAATAACTAATTCCTC<br>CTACGTTCACCCTCTAAAGGTAACAGTTATCTTTGTGGTT<br>CTGAGCCTCCGGAGACTTGGGACGAACTTCACAGCCCTG<br>GGTGTGTGGGCTCTATGCTTTCTCCTGAAATGTGGTTCAT<br>GGTTAGCTTGGGTTTGGGTTGCTACTTTGAAGAAATAAAG<br>CAGGCTGGGTATGGTAGCACAGGCCTTTAAACCCAACAT<br>TCCAAAGGCTGAGGCAGACAGATGCATCTCTGGGAGGAG<br>GAGGCCAGCCTGGTAACACAGTTAAATAAATCCTGGCGC<br>ACACACACAGAACTATTTACTACATTGCGGAGAGAGTAG<br>AGGCTACGGAAGACCTGAAGTACCACCTGCCCTTGTTTTG<br>GGACAAGACGTCACACTGTATCCCAGGATGGCCTGGAAT<br>TCACTATGTAAACTAGGTTGACTTTGAATTTGCAATGATC<br>TTCCTTCCTCTACCTCCTGCCAAGATTATTAGGCATGGCC<br>CACCACACCTGGCTCACTCATTTTAGTATCCTGGAGTATT<br>GTACCGCATACAAACCCTAATATATATATATAATATTTGGTT<br>GCAGGAGGTGGGTTGTGGGCCCATGTGTGAGAAGACAAC<br>TTGTCGGGGCTAGTTCTCTCCCTCAGCCATGTGGATTCTG<br>GGGCCTGAACCCAGGTCTGCAAGCAAGTGGCCAGCATCT<br>TTCCTTGCAGACTCTGCCGCACACTGATTTTTTGAGGGG<br>GGTGGGGTGGGGTGAGACGGGGGGGAAGGACCTTGATAT<br>ATAGTTCTCTACTGGCTTAGAACACCCTTGTGTCAGGAAT<br>AGTGACACTCACGCCTTTAAACCTAGCACTCGGGAGGCA<br>GAGGCAGGCAGATTTCTGAGTTTGAGGCCAGCCTGGTCC<br>ATAGAGTGAGTTCCAGGACAGCCAGGGCTATACAGAGAA<br>ACCCTGTCTCAAAAAAAAAAAAAAAAAAAAAAAACTCCTT<br>CTATACATCAGGCTGGCTCCAACTTGCAGAGATCTGCTTG<br>CCTCTGCCCTCTGCGCGAGTGCTGGAACTACAGGCATGCA<br>CAACACGCCCAGCCTCAGACTCAGCCTCTTAATCAAATGC<br>ACAGAAATAAGTTGGATGCTTCTCTTGAGATGTTGCCTAA<br>GTATATATGGATATACTTACTCATTTATTGCTATTGGGAA<br>AGACTCTGCTCTGGTACAGTTACAGTGACTCACAATTATA<br>ATTCCAGTACTGAGGAGTATGAGGGAAAGGAATTGCTTC<br>AGGTAAGAAGTTACCCTGGGCTATACAACAAGATTCCAT<br>CTTGAACCTCCCTCTTTTACACTGAAAGGAAGTCCGGGGT<br>GATTGCTAGCAGGAAACTTTTCTGATTGAAAGACAGCCC<br>AAAGCTAGTAAGTCACACACTTCAGTCACGGGTGCAAAG<br>TCCCCCAGGTAACTGACACTGGTCACTTCTTAGGAACCAT<br>TGAGTCAGGGAAAGAATCAATCTGAGTGTATACATATAG<br>CATACGTCA\|TACGTGGTTAGAGCACTTACTGCTTTTTCCA<br>GAGCTGACTGTAACTCTAGCCAGGGGATGTGTTGTCCTCT<br>TCTGGCCTCTGCCTACAGCAAGTGCCCACATATATGTGCA<br>GGTATGTATATGATGGCATATGATCCCAGCAATCTTCCAC<br>TCAGAGGCAGCAGAAGCCCTGTCTAAAGAAAGACTGGTA<br>CAGGAAAATGCAGAGTTAATGTAATGGGGTGGAGGAACA<br>AGAGTCGGGGGCGGGGGGGGGGGGGGGGTATCTCAGG<br>CCTGAAGTCCCAAGAATCAAGAAGCTGAGGCAGGAGAAT<br>AGTCTCCAGCTTGAGGTCAGCCCAGGTTAAAAATGACAG |

<table>
<tr><td></td><td></td><td></td><td>CTTGTAGAGAATGGGAGAAGAAGAGAAATTAATGGTTGG<br>TTAATATGATCAC|GTTCGAGAACAGTCAGGGATGCACAG<br>AGAAACCGTCTCAAAAGACAAATACAGTAACCAAGACCA<br>AAACCAACAACCAAACCAAACCAAACACAAACCCCAAA<br>ACAAAAGGCTAAGCTATTCCTCCTAACTCTTAGCATTGCT<br>CAAAGTGTGCCTGCTGTCCTAGGGGCAGGCCCTGGGCCA<br>GATTCACCCCGGTTAAGCAGAATATATGTCAGGTCTCCTG<br>TGGGCCAGAGATTTTGCAGATGAGGTGTTTTTGACTCTAA<br>CAGTATAGTACCAGACTGTCATCTCAGCACTTAGGAGGCT<br>GAGGCAGGGGGATTGCAATGAGTTCCAGGTTAGTTCTCTT<br>CAGTGAGACCCTGATTCAAAACCAAAACAAGGCCAAGGT<br>TGAGCTCTAGTTGGCAGAATGCTCATCAGGTAGTTCTTGG<br>AGGCTTTGAATCTGGAATCCCCTAGCCTCAGCCTTCCAGG<br>AGCTAGATTACAAGCAAGTGCTGTTTCTATTAAATACCAT<br>GTTTTGGGTTTGGGGAGATGGGCCAGTGGTCAAGAGTGC<br>TTGCTTCTCCTTCAGTGGACCAGACAGACTTCAGTTCCCA<br>GCATCCAGGTTGGGTGGTCCACAGCCCCCTAACATTCCAG<br>CTTCAGGGGAATCTGATACCCTCT</td></tr>
<tr><td>ΔSRR107-111/+<br>chr3:34757618-<br>34762637</td><td>A2</td><td>Y</td><td>CCCTGAGCTGGATGTTAGGGGGCTGTGGACCACCCAACC<br>TGGATGCTGGGAACTGAAGTCTGTCTGGTCCACTGAAGG<br>AGAAGCAAGCACTCTTGACCACTGGCCCATCTCCCCAAA<br>CCCAAAACATGGTATTTAATAGAAACAGCACTTGCTTGTA<br>ATCTAGCTCCTAGAAGGCTGGGGCTAGGGGATTCCAGAT<br>TCAAAGCCTCCAAGAACTACCTGATGAGCATTCT|ATGGG<br>GGACTTTTGGTATAGCATTGGAAATGTAAATGAGCTAAA<br>TACCTAATAAAAAATGAAAAAAAAATGTACGTTTAAACT<br>CAAAATCATGATGTCATGATGATGAAGTGCTGGGGGAGC<br>AAGACAGGGCCATTGCCAGGGTAGCCTGTGCTACAGAAT<br>AAGACCCGGTCTCAAAAGAGAGCGCCGAGGGTGGGGGA<br>ACTTCCCTACGCCATCATCCCCCCCACACCCCTCCCCCAA<br>AGAGGGGGAAAAGGCAAAAAAGCAGTAAATAAGAGATA<br>GTCTAGTGGTACCGCCTGTCATCCCAGCTACTCAGGAAGC<br>TAAGGCAGTTACCTGATGAGTGTAAGGCCTGCCTGGGCT<br>ACATGGGTTCAAAGCTAACCCTGGGCAACTTACTGAAAC<br>TCTTTCAAAAGATAAAAAAGAGAACTGG</td></tr>
<tr><td rowspan="2">ΔSRR107/+<br>chr3:34757618-<br>34759104</td><td>C2</td><td>N</td><td>AATAATTGAGGCCATGCTAGTCTACAGATTGAGTTCCAG<br>GACAGGCCGGGATGCACAGAGAAACCCTGTCTTGAAAAC<br>ACCCCCAAAAAATCAATCCAGTAGATGGAATAAAGTATT<br>TTGTGAATGATCTCAAACTCCTGATTGTCAAGGTTCCCCT<br>GGCATGAATGGTCTTTATTGTTAATAATAGTTCTTCTCGT<br>GATCAAATATATACCTAAATGATCTTAGTTCAATGTATCA<br>TTCCTTGGCCAGATG|GGGTGGGTATGGGGGACTTTTGGT<br>ATAGCATTGGAAATGTAAATGAGCTAAATACCTAATAAA<br>AAATGAAAAAAAAATGTACGTTTAAACTCAAAATCATGA<br>TGTCATGATGATGAAGTGCTGGGGGAGCAGAGACAGGGC<br>CATTGCCAGGGTAGCCTGTGCTACAGAATAAGACCCGG</td></tr>
<tr><td>C8</td><td>N</td><td>ACTGGATATTAAATAATTGAGGCCATGCTAGTCTACAGAT<br>TGAGTTCCAGGACAGGCCGGGATGCACAGAGAAACCCTG<br>TCTTGAAAACACCCCCAAAAAATCAATCCAGTAGATGGA<br>ATAAAGTATTTTGTGAATGATCTCAAACTCCTGATTGTCA<br>AGGTTCCCCTGGCATGAATGGTCTTTATTGTTAATAATAG<br>TTCTTCTCGTGATCAAATATATACCTAAATGATCTTAGTT<br>CAATGTATCATTCCTTGGCCAGATG|GTATAGCATTGGAA<br>ATGTAAATGAGCTAAATACCTAATAAAAAATGAAAAAAA<br>AATGTACGTTTAAACTCAAAATCATGATGTCATGATGATG</td></tr>
</table>

| | | | |
|---|---|---|---|
| | | | AAGTGCTGGGGGAGCAGAGACAGGGCCATTGCCAGGGTA GCCTGTGCTACAGAATAAGACCCGG |
| | G5 | N | GCACTGGATATTAAATAATTGAGGCCATGCTAGTCTACA GATTGAGTTCCAGGACAGGCCGGGATGCACAGAGAAACC CTGTCTTGAAAACACCCCCAAAAAATCAATCCAGTAGAT GGAATAAAGTATTTTGTGAATGATCTCAAACTCCTGATTG TCAAGGTTCCCCTGGCATGAATGGTCTTTATTGTTAATAA TAGTTCTTCTCGTGATCAAATATATACCTAAATGATCTTA GTTCAATGTATCATTCCTTGGCCAGATG\|TGGGTATGGGG GACTTTTGGTATAGCATTGGAAATGTAAATGAGCTAAAT ACCTAATAAAAAATGAAAAAAAAATGTACGTTTAAACTC AAAATCATGATGTCATGATGATGAAGTGCTGGGGGAGCA GAGACAGGGCCATTGCCAGGGTAGCCTGTGCTACAGAAT AAGA |
| ΔSRR109/+ chr:34759104-34760865 | 34o2 | Y | AAATAAAAGATTAAGCAGTGTGATATACAAGCCTCTAAG ACACACATAAAATGAACCTTATTGCTTATTGAAACTGGGT CTTATTATACAGCCAAGGCTAGCCCAGCACAAAATAAAA TTTAAGATGGAAATT\|GGCTCAAACCTGTAATCCCGACTC TGGAAGCTGATTGTTT |
| ΔSRR111/+ chr:34760865-34762618 | D2 | N | ATATGTCTTATTATTTATTTTATGAGTATGAATATTTT GCCTGTTTGTATGTCTGTGCACCATGTGCATGCCTGGT GCTAATGGAGGCCAGAGGAGGGCATCAGGCCCTCTGG AGCTAGAGTTACAGATGGTTGTGAGCCTCT\|AAAACCAAA ACAAGGCCAAGGTTGAGCTCTAGTTGGCAGAATGCTCAT CAGGTAGTTCTTGGAGGCTTTGAATCTGGAATCCCCTAGC CCCAGCCTTCTAGGAGCTAGATTACAAGCAAGTGCTGTTT CTATTAAATACCATGTTTTGGGTTTGGGGAGATGGGCCAG TGGTCAAGAGTGCTTGCTTCTCCTTCTAGTGGACCAGACA GACT |
| | D9 | N | ATGAATATTTTGCCTGTTTGTATGTCTGTGCACCATGT GCATGCCTGGTGCTAATGGAGGCCAGAGGAGGGCATCAG GCCCTCTGGAGCTAGAGTTACAGATGGTTGTGAGCCTCTA TGCACATGCTGG\|GCAGAATGCTCATCAGGTAGTTCTTGG AGGCTTTGAATCTGGAATCCCCTAGCCCCAGCCTTCTAGG AGCTAGATTACAAGCAAGTGCTGTTTCTATTAAATACCAT GTTTTGGGTTTGGGGAGATGGGCCAGTGGTCAAGAGTGC TTGCTTCTCCTTCAGTGGACCA |
| | G1 | N | AATATTTTGCCTGTTTGTATGTCTGTGCACCATGTGCATG CCTGGTGCTAATGGAGGCCAGAGGAGGGCATCAGGCCCT CTGGAGCTAGAGTTACAGATGGTTGTGAGCCTCTA\|AACC AAGTTGGCAG\|AATGCTCATCAGGTAGTTCTTGGAGGCTT TGAATCTGGAATCCCCTAGCCCCAGCCTTCTAGGAGCTAG ATTACAAGCAAGTGCTGTTTCTATTAAATACCATGTTTTG GGTTTGGGGAGATGGGCCAGTGGTCAAGAGTGCTTGCTT CTCCTTCAGTGGACCAGACA |
| ΔSRR107+111/+ chr3:34757618-34759104 + chr:34760865-34762618 | A5 | Y | GAGTATGATATTTTGCCTGTTTGTATGTCTGTGCACCATG TGCATGCCTGGTGCTAATGGAGGCCAGAGGAGGGCATCA GGCCCTCTGGAGCTAGAGTTACAATGGTTGTGAGCCTCTA \|TTGGCAGAATGCTCATCAGGTATTCTTGGAGGCTTTGAAT CTGGAATCCCCTACCCCACCTTCTAGGACTAATTACAACA ATGCTGTTTCTATTAAATACCATGTTTTGG |
| | A7 | N | ATTTTGCCTGTTTGTATGTCTGTGCACCATGTGCATGCCTG GTGCTAATGGAGGCCAGAGGAGGGCATCAGGCCCTCTGG AGCTAGAGTTACAGATG\|AAAACCAAAACAAGGCCAAGG TTGAGCTCTAGTTGGCAGAATGCTCATCAGGTAGTTCTTG |

| | | | |
|---|---|---|---|
| | | | GAGGCTTTGAATCTGGAATCCCCTAGCCCCAGCCTTCTAG<br>GAGCTAGATTACAAGCAAGTGCTGTTTCTATTAAATACCA<br>TGTTTTGGGTTTGGGGAGATGGGCCAGTGGTCAAGAGTG<br>CTTGCTTCTCCTTCAGTGGACCAGACAGACTTCAGTTC |
| | A9 | N | ATTTTGCCTGTTTGTATGTCTGTGCACCATGTGCATGCCTG<br>GTGCTAATGGAGGCCAGAGGAGGGCATCAGGCCCTCTGG<br>AGCTAGAGTTACAGATGGTTGTGAGCCTCT\|GTTGGCAGA<br>ATGCTCATCAGGTAGTTCTTGGAGGCTTTGAATCTGGAAT<br>CCCCTAGCCCCAGCCTTCTAGGAGCTAGATTACAAGCAA<br>GTGCTGTTTCTATTAAATACCATGTTTTGGGTTTGGGGAG<br>ATGGGCCAGTGGTCAAGAGTGCTTGCTTCTCCTTCAGTGG<br>ACCAGACAGACTTCAGTTC |
| | B4 | N | TGAATATTTTGCCTGTTTGTATGTCTGTGCACCATGTGCAT<br>GCCTGGTGCTAATGGAGGCCAGAGGAGGGCATCAGGCCC<br>TCTGGAGCTAGAGTTACAGATGGTTGTGAGCCTCT\|AAAA<br>CCAAAACAAGGCCAAGGTTGAGCTCTAGTTGGCAGAATG<br>CTCATCAGGTAGTTCTTGGAGGCTTTGAATCTGGAATCCC<br>CTAGCCCCAGCCTTCTAGGAGCTAGATTACAAGCAAGTG<br>CTGTTTCTATTAAATACCATGTTTTGGGTTTGGGGAGATG<br>GGCCAGTGGTCAAGAGTGCTTGCTTCTCCTTCA |
| ΔOS_SRR107+111<br>/+ (OS intact)<br>chr3:34758043-<br>34758115<br><br>the intact OS motif<br>is marked in bold<br>with an underline | A2 | N | CTTCTGGGTGGTGAACCTTGGCA\|CATAATGGGGCT**AAAT<br>AAATAACAATG**\|GGACTATGCTAACCTTCCTGGGTAACAG<br>CCGGGAGGGAGGTGTCATT |
| | E2 | N | CTTCTGGGTG\|GGGGC**TAATAAATAACAATG**ACAGTACTT<br>GCCCTTAGCAGAGTCCTGGGACTATGCTAAACAACTTCCT<br>GGGTAACAGCCGGGAGGGAGGTGTC |
| | H8 | N | CTTCTGGGTGGTGAA\|CCTTGGCA\|TATTATTTTAGC\|TAA<br>TGGGGC**TAA\|TAAATAACAATG**A\|ACTTGCC\|TTAC\|AGTCC<br>TGGGACTATGCTAA\|CAACTTCCTGGG\|AC\|GCCGGGAGGG<br>AGGTGTCATT |
| ΔOS_SRR107+111<br>/+<br>chr3:34758043-<br>34758115 | B1 | N | CTTCTGGGTGGTGAACCTTGGC\|TATGCTAAACAACTTCCT<br>GGGTAACAGCCGGGAGGGAGGTGTCATT |
| | C2 | N | CTTCTGGGTGGTGAACCTTGGCA\|CCTGGGACTATGCTAA<br>ACAACTTCCTGGGTAACAGCCGGGAGGGAGGTGTCATT |
| | F5 | N | CTTCTGGGTGGTGAACCTTGG\|CAGAGTCCTGGGACTATG<br>CTAAACAACTTCCTGGGTAACAGCCGGGAGGGAGGTGTC<br>ATT |
| ΔSRR107+<br>K(2)_111/+<br>chr3:34761207-<br>34761274 | B1 | N | AGCCAGAGATAACCTGGTGGTTGAAGGCAGCCTTC\|CCAG<br>GGTGCCAACTTTGAAGGGCCACAGTAAAGATTAAATTGT<br>ATGTCCCACCTTTATAGCACTCAGGGGGCTGAGGCAGGA<br>GCATCAGGAATCCGAGGCCTTAGCTACGAAACAGGTTCG<br>AGACCAGCTGCAGTTACAAGAAACCCTCTCTCAATTTCAA<br>TGTCCTGTACCCCACCAA |
| | B11 | N | TCACCTTGAGCCAGAGATAACCTGGT\|TAGG\|TCATCTCAT<br>CTTCTAAACCATCCCAGGGTGCCAACTTTGAAGGGCCAC<br>AGTAAAGATTAAATTGTATGTCCCACCTTTATAGCACTCA<br>GGGGGCTGAGGCAGGAGCATCAGGAATCCGAGGCCTTAG<br>CTACGAAACAGGTTCGAGACCAGCTGCAGTTACAAGAAA<br>CCCTCTCTCAATTTCAATGTCCTGTACCCCACCAA |
| | D1 | N | GTCACCTTGAGCCAGAGATAACCTGGTGGTTGAAGGCAG<br>CCTTC\|ATCTCATCTTCTAAACCATCCCAGGGTGCCAACTT<br>TGAAGGGCCACAGTAAAGATTAAATTGTATGTCCCACCTT<br>TATAGCACTCAGGGGGCTGAGGCAGGAGCATCAGGAATC |

| | | | |
|---|---|---|---|
| | | | CGAGGCCTTAGCTACGAAACAGGTTCGAGACCAGCTGCA<br>GTTACAAGAAACCCTCTCTCAATTTCAATGTCCTGTACCC<br>CACCAA |
| | G2 | N | CCTTGAGCCAGAGATAACCTG\|CCTTTCATCTCATCTTCTA<br>AACCATCCCAGGGTGCCAACTTTGAAGGGCCACAGTAAA<br>GATTAAATTGTATGTCCCACCTTTATAGCACTCAGGGGGC<br>TGAGGCAGGAGCATCAGGAATCCGAGGCCTTAGCTACGA<br>AACAGGTTCGAGACCAGCTGCAGTTACAAGAAACCCTCT<br>CTCAATTTCAATGTCCTGTACCCCACCAA |
| | H9 | N | GAGGGTTCTTGTACTGCAGCTGGTCTCGAACCTGTTTCGT<br>AGCTAAGGCCTCGGATTCCTGATGCTCCTGCCTCAGCCCC<br>CTGAGTGCTATAAAGGTGGGACATACAATTTAATCTTTAC<br>TGTGGCCCTTCAAAGTTGGCACCCTGGGATGGTTTAGAAG<br>ATGAGATGAAA\|CCTCGTTAATAGAAGAATTTAAGAATGA<br>CTCAAATGGAAGGTGGAGGACAATTAGGGTTTAAAAAAA<br>GAACCTGGGATGGGCCAGTTGTAAACCCCCTGGAGCTGC<br>CTAGAGGAAGGAGCTGGAGGAGAGCTTAGAAAACAAAG<br>GGGGAGGTCATGGAAACAGACGGGGAGGTCAGACA |
| ΔSRR85-95/+<br>chr3:34733021-<br>34748441 | E2 | N | CTGCATGGAAGTTCCTAGACCAGTGTCTGGTGCTCTGGAG<br>TGAGTGATGTCACTGGGTTCTGGATATCAGGTGCAGCCA\|<br>TGCGTGTCATACTGTTTTAAGATCAGAAATGCTAAAGGTT<br>CAGTCAATTTTCATGGTTCTACTTTGACACTCTCCCGCAG<br>AACTTATGGTCTGTTTAAAAATAGAAAACGCAGCCATCT<br>GGCTATTTGATGGGATTCTATTTTTGTTTCTCTTTGCGTTC<br>GCAAAGTGTGTTGGGTCTGAAATTTTCCGTGTTCTGCCCT<br>ATATGTAATTGTGTGTATATACACACATACTTTCTCATTT<br>AAATCTCCATACACTTCCTCATCTAAATCTTGCTGTTATC<br>AGTCTGTGTTGTTTGTGGTCCACGGCAGTGTTTGGTCGGG<br>ATGTCAACCTTGCTTAGTTCATCCACTAGTCACGTCTGCA<br>CTGAATTCCTACTCTAAATTCTTACCAAA |
| | F6 | N | TGCATGGAGTTCTAGACCAGTGTCTGGTGCTCTGGAGTGA<br>GTGATGTCACTGGGTTCTGGATATCAGGTGCAGCCATGG\|<br>CACTGCGTGTCATACTGTTTTAAGATCAGAAATGCTAAAG<br>GTTCAGTCAATTTTCATGGTTCTACTTTGACACTCTCCCGC<br>AGAACTTATGGTCTGTTTAAAAATAGAAAACGCAGCCAT<br>CTGGCTATTTGATGGGATTCTATTTTTGTTTCTCTTTGCGT<br>TCGCAAAGTGTGTTGGGTCTGAAATTTTCCGTGTTCTGCC<br>CTATATGTAATTGTGTGTATATACACACATACTTTCTCATT<br>TAAATCTCCATACACTTCCTCATCTAAATCTTGCTGTTATC<br>AGTCTGTGTTGTTTGTGGTCCACGGCAGTGTTTGGTCGGG<br>ATGTCAACCTTGCTTAGTTCATCCACTAGTCACGTCTGCA<br>CTGAATTCCTACTCTAAATTCTTACCAAAGGTCCCC |
| | G4 | N | TGTCAAGTGGACTGCATGGAGTTCCTAGACCAGTGTCTGG<br>TGCTCTGGAGTGAGTGATGTCACTGGGTTCTGGATATCAG<br>GTGCAGCCATGGG\|CACTGCGTGTCATACTGTTTTAAGAT<br>CAGAAATGCTAAAGGTTCAGTCAATTTTCATGGTTCTACT<br>TTGACACTCTCCCGCAGAACTTATGGTCTGTTTAAAAATA<br>GAAAACGCAGCCATCTGGCTATTTGATGGGATTCTATTTT<br>TGTTTCTCTTTGCGTTCGCAAAGTGTGTTGGGTCTGAAAT<br>TTTCCGTGTTCTGCCCTATATGTAATTGTGTGTATATACAC<br>ACATACTTTCTCATTTAAATCTCCATACACTTCCTCATCTA<br>AATCTTGCTGTTATCAGTCTGTGTTGTTTGTGGTCCACGG<br>CAGTGTTTGGTCGGGATGTCAACCTTGCTTAGTTCATCCA<br>CTAGTCACGTCTGCACTGAATTCCTACTCTAAATTCTTAC<br>CA |

| | | | |
|---|---|---|---|
| ΔSRR104-<br>dCTCF/+<br>chr3:34755000-<br>34774122 | B6 | Y | GTAGTGACTGCAGCAGACTTGGGAAGATACTTTACCATC<br>CCACAGCTGAGAGCCACTGAGACCGAGGTTTAGAATTTC<br>ATCCTCAAGCCAAGATACTAAACATATCAATGAATGCGG<br>ATGCCTTGCTATGCCCAGAATTCCCTCTCCGTCTCCAAGC<br>CTTACGGGAACGCCATATGCCAGGGGTTCCTGGCAGCAG<br>GAAACCAAGAGACTAACAGAATAAATTACTTTACATTAG<br>ACACGTGCTGTTGACCTGCTCGAGGTATGAAG\|TGGTTAG<br>AGCACTTACTGCTTTTCCAGGGGACCTGGGATGGCTCCTC<br>CCCACCCACATGGTGGTTCAGAGCTGACTGTAACTCTAGC<br>CAGGGGATGTGTTGTCCTCTTCTGGCCTCTGCCTACAGCA<br>AGTGCCCACATATATGTGCAGGTATGTATATGATGGCATA<br>TGATCCCAGCAATC |
| | B8 | N | TCCAAGCCAAGGCTCAGCGACTCTGAGTCCCAACATCAC<br>TGTAGTGACTGCAGCAGACTTGGGAAGATACTTTACCATC<br>CCACAGCTGAGAGCCACTGAGACCGAGGTTTAGAATTTC<br>ATCCTCAAGCCAAGATACTAAACATATCAATGAATGCGG<br>ATGCCTTGCTATGCCCAGAATTCCCTCTCCGTCTCCAAGC<br>CTTACGGGAACGCCATATGCCAGGGGTTCCTGGCAGCAG<br>GAAACCAAGAGACTAACAGAATAAATTACTTTACATTAG<br>ACACGTGCTGTTGACCTGCTCGAGGTATGAAGAATATTA<br>ACACCGTCCCCG\|GTAGCTCAGTGGTTAGAGCACTTACTG<br>CTTTTCCAGGGGACCTGGGATGGCTCCTCCCCACCCACAT<br>GGTGGTTCAGAGCTGACTGTAACTCTAGCCAGGGGATGT<br>GTTGTCCTCTTCTGGCCTCTGCCTACAGCAAGTGCCCACA<br>TATATGTGCAGGTATGTATATGATGGCATATGATCCCAGC<br>AATC |
| | E6 | N | CCAAGGCTCAGCGACTCTGAGTCCCAACATCACTGTAGT<br>GACTGCAGCAGACTTGGGAAGATACTTTACCATCCCACA<br>GCTGAGAGCCACTGAGACCGAGGTTTAGAATTTCATCCTC<br>AAGCCAAGATACTAAACATATCAATGAATGCGGATGCCT<br>TGCTATGCCCAGAATTCCCTCTCCGTCTCCAAGCCTTACG<br>GGAACGCCATATGCCAGGGGTTCCTGGCAGCAGGAAACC<br>AAGAGACTAACAGAATAAATTACTTTACATTAGACACGT<br>GCTGTTGACCTGCTCGAGGTATGAAGAATATTAACACCGT<br>CCCCGAACGG\|TAGCTCAGTGGTTAGAGCACTTACTGCTT<br>TTCCAGGGGACCTGGGATGGCTCCTCCCCACCCACATGGT<br>GGTTCAGAGCTGACTGTAACTCTAGCCAGGGGATGTGTT<br>GTCCTCTTCTGGCCTCTGCCTACAGCAAGTGCCCACATAT<br>ATGTGCAGGTATGTATATGATGGCATATGATCCCAGCAA |
| ΔSRR85-95+SCR-<br>dCTCF/+<br>chr3:34733021-<br>34748441<br>+<br>chr3:34755000-<br>34774122 | B7 | N | CAAGCCAAGGGCTCAGCGACTCTGAGTCCCAACATCACT<br>GTAGTGACTGCAGCAGACTTGGGAAGATACTTTACCATC<br>CCACAGCTGAGAGCCACTGAGACCGAGGTTTAGAATTTC<br>ATCCTCAAGCCAAGATACTAAACATATCAATGAATGCGG<br>ATGCCTTGCTATGCCCAGAATTCCCTCTCCGTCTCCAAGC<br>CTTACGGGAACGCCATATGCCAGGGGTTCCTGGCAGCAG<br>GAAACCAAGAGACTAACAGAATAAATTACTTTACATTAG<br>ACACGTGCTGTTGACCTGCTCGAGGTATGAAGAATATTA<br>ACACCGTCCCCGAACGG\|TAGCTCAGTGGTTAGAGCACTT<br>ACTGCTTTTCCAGGGGACCTGGGATGGCTCCTCCCCACCC<br>ACATGGTGGTTCAGAGCTGACTGTAACTCTAGCCAGGGG<br>ATGTGTTGTCCTCTTCTGGCCTCTGCCTACAGCAAGTGCC<br>CACATATATGTGCAGGTATGTATATGATGGCATATG |
| | C11 | Y | GGAAGATACTTTACCATCCCACAGCTGAGAGCCACTGAG<br>ACCGAGGTTTAGAATTTCATCCTCAAGCCAAGATACTAA<br>ACATATCAATGAATGCGGATGCCTTGCTATGCCCAGAATT<br>CCCTCTCCGTCTCCAAGCCTTACGGGAACGCCATATGCCA |

<table>
<tr><td></td><td></td><td></td><td>GGGGTTCCTGGCAGCAGGAAACCAAGAGACTAACAGAAT<br>AAATTACTTTACATTAGACACGTGCTGTTGACCTGCTCGA<br>GGTATGAAGAATATTAACACCGTCCCCGAACGG|TAGCTC<br>AGTGGTTAGAGCACTTACTGCTTTTCCAGGGGACCTGGGA<br>TGGCTCCTCCCCACCCACATGGTGGTTCAGAGCTGACTGT<br>AACTCTAGCCAGGGGATGTGTTGTCCTCTTCTGGCCTCTG<br>CCTACAGCAAGTGCCCACATATATGTGCAGGTATGTATAT<br>GATGGCATATGATC</td></tr>
<tr><td></td><td>D2</td><td>N</td><td>GTAGTGACTGCAGCAGACTTGGGAAGATACTTTACCATC<br>CCACAGCTGAGAGCCACTGAGACCGAGGTTTAGAATTTC<br>ATCCTCAAGCCAAGATACTAAACATATCAATGAATGCGG<br>ATGCCTTGCTATGCCCAGAATTCCCTCTCCGTCTCCAAGC<br>CTTACGGGAACGCCATATGCCAGGGGTTCCTGGCAGCAG<br>GAAACCAAGAGACTAACAGAATAAATTACTTTACATTAG<br>ACACGTGCTGTTGACCTGCTCGAGGTATGAAGAAT|AACG<br>GTAGCTCAGTGGTTAGAGCACTTACTGCTTTTCCAGGGGA<br>CCTGGGATGGCTCCTCCCCACCCACATGGTGGTTCAGAGC<br>TGACTGTAACTCTAGCCAGGGGATGTGTTGTCCTCTTCTG<br>GCCTCTGCCTACAGCAAGTGCCCACATATATGTGCAGGTA<br>TGTATATGATGGCATATGAT</td></tr>
<tr><td>ΔSRR85-95+107-dCTCF/+<br>chr3:34733021-34748441<br>+<br>chr3:34757641-34774122</td><td>A12</td><td>Y</td><td>TAAAGTTTAAACGTACATTTTTTTTTCATTTTTTATTAGGT<br>ATTTAGCTCATTTACATTTCCAATGCTATACCAAAAGTCC<br>CCCATACCCACCCAACG|TCGGGGACGGTGTTAATATTCT<br>TCATACCTCGAGCAGGTCAACAGCACGTGTCTAATGTAA<br>AGTAATTTATTCTGTTAGTCTCTTGGTTTCCTGCTGCCAGG<br>AACCCCTGGCATATGGCGTTCCCGTAAGGCTTGGAGACG<br>GAGAGGGAATTCTGGGCATAGCAAGGCATCCGCATTCAT<br>TGATATGTTTAGTATCTTGGCTTGAGGATGAAATTCTAAA<br>CCTCGGTCTCAGTGGCTCTCAGCTGTGGGATGGTAAAGTA<br>TCTTCCCAAGTCTGCTGCAGTCACTACAGTGATGTTGGGA<br>CTCAGAGTCGCTGAGCCTTGGCTTGGAGACCTGATAAGG<br>GCTTGTAAGAGTAGTACCTCAGTCTCCCTAAGGCCTGCCT<br>GGAGTTCTGCACTGCAACTGTGTCCGAGGAGTCCTCCCTT<br>AA</td></tr>
<tr><td></td><td>C3</td><td>N</td><td>CAATTCATCATCAAGACATCATGATTTTGAGTTTAAACGT<br>ACATTTTTTTTTCATTTTTTATTAGGTATTTAGCTCATTTA<br>CATTTCCAATGCTATACCAAAAGTCCCCCATACCCACCCA<br>ACG|TCGGGGACGGTGTTAATATTCTTCATACCTCGAGCA<br>GGTCAACAGCACGTGTCTAATGTAAAGTAATTTATTCTGT<br>TAGTCTCTTGGTTTCCTGCTGCCAGGAACCCCTGGCATAT<br>GGCGTTCCCGTAAGGCTTGGAGACGGAGAGGGAATTCTG<br>GGCATAGCAAGGCATCCGCATTCATTGATATGTTTAGTAT<br>CTTGGCTTGAGGATGAAATTCTAAACCTCGGTCTCAGTGG<br>CTCTCAGCTGTGGGATGGTAAAGTATCTTCCCAAGTCTGC<br>TGCAGTCACTACAGTGATGTTGGGACTCAGAGTCGCTGA<br>GCCTTGGCTTGGAGACCTGATAAGGGCTTGTAAGAGTAG<br>TACCTCAGTCTCCCTAAGGCCTGCCTGGAGTTCTGCAAAA<br>AAACTGTGTCAAAAGAAAAAC</td></tr>
<tr><td></td><td>D12</td><td>N</td><td>AATTCATCATCAAGACATCATGATTTTGAGTTTAAACGTA<br>CATTTTTTTTTCATTTTTTATTAGGTATTTAGCTCATTTAC<br>ATTTCCAATGCTATACCAAAAGTCCCCCATACCCACCCAA<br>CG|TCGGGGACGGTGTTAATATTCTTCATACCTCGAGCAG<br>GTCAACAGCACGTGTCTAATGTAAAGTAATTTATTCTGTT<br>AGTCTCTTGGTTTCCTGCTGCCAGGAACCCCTGGCATATG<br>GCGTTCCCGTAAGGCTTGGAGACGGAGAGGGAATTCTGG<br>GCATAGCAAGGCATCCGCATTCATTGATATGTTTAGTATC</td></tr>
</table>

<table>
<tr><td></td><td></td><td></td><td>TTGGCTTGAGGATGAAATTCTAAACCTCGGTCTCAGTGGC<br>TCTCAGCTGTGGGATGGTAAAGTATCTTCCCAAGTCTGCT<br>GCAGTCACTACAGTGATGTTGGGACTCAGAGTCGCTGAG<br>CCTTGGCTTGGAGACCTGATAAGGGCTTGTAAGAGTAGT<br>ACCTCAGTCTCCCTAAGGCCTGCCTGGAGTTCTGCACAGC<br>AACTGTGTCCAAGGA</td></tr>
<tr><td></td><td>H1</td><td>N</td><td>TTCTTCATCAAGACATCATGATTTTGAGTTTAAACGTACA<br>TTTTTTTTTCATTTTTTATTAGGTATTTAGCTCATTTACATT<br>TCCAATGCTATACCAAAAGTCCCCCATACCCACCCAACG|<br>TCGGGGACGGTGTTAATATTCTTCATACCTCGAGCAGGTC<br>AACAGCACGTGTCTAATGTAAAGTAATTTATTCTGTTAGT<br>CTCTTGGTTTCCTGCTGCCAGGAACCCCTGGCATATGGCG<br>TTCCCGTAAGGCTTGGAGACGGAGAGGGAATTCTGGGCA<br>TAGCAAGGCATCCGCATTCATTGATATGTTTAGTATCTTG<br>GCTTGAGGATGAAATTCTAAACCTCGGTCTCAGTGGCTCT<br>CAGCTGTGGGATGGTAAAGTATCTTCCCAAGTCTGCTGCA<br>GTCACTACAGTGATGTTGGGACTCAGAGTCGCTGAGCCTT<br>GGCTTGGAGACCTGATAAGGGCTTGTAAGAGTAGTACCT<br>CAGTCTCCCTAAGGCCTGCCTGGAGTTCTGCACTGCAACT<br>GTG</td></tr>
</table>

**Supplementary Table S4: Guide sequences for insertion lines**

| Name | Target site | Sequence |
|---|---|---|
| *Sox2* | 3’ coding sequence of *Sox2* | CCCCTGTCGCACATGTGA |
| MH5’ | 5’ of P2A-Venus cassette | TTCCTCCCATGTGCGCCC |
| MH3’ | 3’ of P2A-Venus cassette | CAAGTAATGAGGGCTCCC |
| Insertion | Intervening region between *Sox2* and SCR | GTTCAAAAACTAGAAACA |

**Supplementary Table S5: qPCR primers for gene expression analysis (SNPs indicated as lowercase)**

| mRNA | Allele | Forward Sequence | Reverse Sequence |
|---|---|---|---|
| Sox2 | 129 | GGACTTCTTTTTGGGGGACT | CGCCTAACGTACCACTAGAACTT**t** |
| Sox2 | CAST | GGACTTCTTTTTGGGGGACT | CGCCTAACGTACCACTAGAACTT**a** |
| Sdha | n/a | ACTGGGATGGGCTCCTTAGT | GCCCTGAGAAAGATCACGTC |
| Gapdh | n/a | GCACCAGCATCCCTAGACC | CTTCTTGTGCAGTGCCAGGTG |

**Supplemental Table S6: 4C primers**

Sequences of 4C primers. Blue denotes Illumina adapter sequence for high-throughput sequencing. Red denotes position of 6-nucleotide barcodes, used to multiplex 4C samples for sequencing.

| Name | Sequence |
|---|---|
| Near-SCR DpnII | 5'-AATGATACGGCGACCACCGAGATCTACACTCTTTCCCTACACGACGCTCTTCCGATCTNNNNNNGCAAGAGCCAGGTGTGGCTC-3' |
| Near-SCR Csp6I | 5'-CAAGCAGAAGACGGCATACGAGCTCTTCCGATCTCCTGGTGCTTTGCCCAGCAC-3' |
| SCR DpnII | 5'-AATGATACGGCGACCACCGAGATCTACACTCTTTCCCTACACGACGCTCTTCCGATCTNNNNNNGGGGAGGTCAGACACCTGATC-3' |
| SCR Csp6I | 5'-CAAGCAGAAGACGGCATACGAGCTCTTCCGATCTTTCCGGTAGGGGTGGAGC-3' |
| hSOX9 DpnII | 5'-AATGATACGGCGACCACCGAGATCTACACTCTTTCCCTACACGACGCTCTTCCGATCTNNNNNNAGGACATTGATTTGGATC-3' |
| hSOX9 Csp6I | 5'-CAAGCAGAAGACGGCATACGAGCTCTTCCGATCTCGTAGTGTGGACCTATTT-3' |
